# Sparse Linear Algebra Accelerates Genotype Representation Graph Computation at Biobank Scale

**DOI:** 10.64898/2026.09.10.750583

**Authors:** Yifan Li, Qingyao Sun, Drew DeHaas, Max Xiaohang Zhao, Adam R. Boyko, Shaila A. Musharoff, Xinzhu Wei, Giulia Guidi

## Abstract

Biobank-scale genomic analyses are increasingly constrained by computational costs, as hundreds of thousands to millions of samples and variants must be analyzed together. The genotype representation graph (GRG) compactly encodes population genetic variation to accelerate computation, but its graph-traversal implementation does not exploit modern accelerators. This work introduces Mikado, a methodology that expresses GRG-based computation using sparse linear algebra primitives. Under a reverse topological ordering, the GRG adjacency matrix is strictly block-lower-triangular, and the genotype matrix-vector product becomes a sparse triangular solve, decomposed into a pipelined sequence of blocked sparse matrix-vector multiplies. Because GRGs are shallow by construction, the solve completes in only a few sequential stages, exposing fine-grained parallelism and enabling vendor-optimized sparse primitives on GPUs. On the full *All of Us* cohort, Mikado computes principal components *in minutes*, where PLINK2 cannot complete one chromosome in 72 hours, and runs genome-wide BOLT-LMM-inf in *under 19 minutes* on one GPU, where the official BOLT-LMM analysis would be impractical. Our two-stage reformulation is independent of the hardware: the same code runs unchanged across GPU architectures and, on CPUs, gives a **1.8**× single-threaded speedup over the existing GRG library together with multi-threaded scaling, providing a scalable, portable approach for population genetics as biobanks continue to grow.

## 1 Introduction

Over the past decade, the scale of population genomics data has increased dramatically. Biobank initiatives such as the *All of Us* Research Program now include hundreds of thousands to a million phased whole genomes [1], along with environmental and phenotypic data. This growth improves statistical power for genome-wide association studies (GWAS), heritability estimation, fine mapping, and emerging applications such as gene–environment interaction (G**×**E) testing [2–7]. The computational demands of these analyses, however, are rapidly outpacing conventional approaches. Common tabular formats such as VCF, BED (PLINK1), and BGEN represent each variant independently and do not exploit the shared relatedness structure underlying population genetic variation [2, 3]. PGEN (PLINK2) can optionally exploit linkage disequilibrium (LD) among nearby single nucleotide polymorphisms (SNPs) to reduce storage [8], but it leaves redundancy in downstream statistical computation unaddressed. In benchmarks reported in concurrent work [9], principal component analysis (PCA) on simulated cohorts of 100k to 1 million individuals (14.5–23.7 million variants) in PGEN format took from several hours to more than 150 hours on a server with two 32-core AMD EPYC 7532 processors and 1 TiB of RAM.

This work introduces Mikado,^1^ a general sparse linear algebra formulation that makes operations on compressed genotype representations efficient and portable on modern accelerators. Its generality rests on a single primitive: repeated matrix–vector multiplication with the genotype matrix, which governs the scalability of many statistical genetics methods and, in some cases, determines their accuracy. In mixed-model association testing, such as in BOLT-LMM [10], this product is the core operation in the inner loop of the conjugate gradient solver. In randomized Haseman–Elston regression [11], multiplying the genotype matrix by random vectors dominates the runtime, and faster trace estimation directly reduces the standard error of the heritability estimates. The same operation underlies randomized PCA, where the top eigenvectors are obtained through iterative methods that repeatedly multiply by the genotype matrix, as well as polygenic risk score computation and LD score regression, which project the genotype matrix onto a precomputed or random vector. Of these, PCA and mixed-model association via BOLT-LMM are the most demanding: both are widely used and statistically standard, and both run many iterations, so the per-product cost compounds and this primitive becomes the limiting factor.

The genotype representation graph (GRG) was recently introduced as a compact substrate for biobank-scale genotype data [6, 9, 12], achieving more than 25**×** compression relative to VCF.GZ on the UK Biobank WGS dataset, which includes 490,541 samples and 706,556,181 variants across 22 chromosomes. A GRG is a directed acyclic graph that captures genetic similarity among samples through shared ancestry-bearing nodes, so highly similar samples reuse a common representation of the variants they share. Because central operations such as allele frequency computation, genotype–phenotype association, and phenotype simulation run directly on the GRG via graph traversal, reusing computed values across shared nodes, they run orders of magnitude faster than equivalent operations on tabular data [6, 9, 12]. As cohorts grow to millions of individuals and analyses expand across many traits, SNPs, and environmental variables, further speed and portability to modern, rapidly evolving hardware become critical.

GPUs deliver much higher throughput and bandwidth than CPUs by using tens of thousands of cores [13], with speedup of more than 100-fold reported for data analytics and machine learning [14–19]. SAIGE-GPU achieves a fivefold speedup over CPU-based SAIGE across more than two thousand phenotypes by porting the existing linear algebra to the GPU without changing the underlying genotype representation [20]. GRG traversal, however, does not map naturally onto GPU execution. The node values must be resolved in topological order. On a single processor, it is sufficient to pick one valid ordering, whereas on a GPU the problem becomes one of scheduling, that is deciding which nodes can be resolved concurrently, at which step, and on which threads, while respecting the dependency structure. The resulting work pattern is irregular, with low-locality memory accesses that degrade load balancing, warp occupancy, and effective memory bandwidth. Realizing the compression advantages of the GRG as GPU throughput therefore requires redesigning the computation rather than simply porting the traversal.

GPU architectures favor dense, regular computation, but sparse computation has also become efficient on them. Prior work has mapped computational biology problems to sparse linear algebra primitives to enable acceleration [16, 17, 21–24]. These primitives, such as sparse matrix–vector multiplication (SpMV) [25, 26], are supported by highly optimized, portable libraries like cuSPARSE [27] and rocSPARSE [28], which hardware vendors and the research community maintain as architectures evolve rapidly. Our key observation is that sparse linear algebra primitives can bridge the gap between GRG traversal and efficient accelerator-based computation.

Methodologically, Mikado reformulates GRG traversal as a blocked sparse triangular solve built from sparse matrix-vector products, using wavefront scheduling that exposes the parallelism accelerators need while preserving GRG compression. The name reflects this scheduling rule, where, as in the game, a block is dispatched only after every block it depends on has been resolved, so computation advances as dependencies are satisfied rather than waiting at a global synchronization barrier. That formulation is implemented as a high-performance, hardware-portable CPU and GPU backend for `grapp` [9], relying only on standard sparse primitives so it carries across accelerator architectures without requiring reimplementation.

Mikado makes analyses that are practically infeasible with standard tools routine on commodity accelerator architectures. On the *All of Us* cohort [1], PCA that PLINK2 cannot complete on a single full chromosome within 72 hours finishes *in minutes*. The mixed-model association with BOLT-LMM-inf runs 26 faster end to end than the official BOLT-LMM distribution on the 1000 Genomes cohort and more than 1,000 faster on a simulated 200,000-individual cohort; on the full *All of Us* cohort, the genome-wide analysis completes under 19 minutes on a single GPU. On the same GPU, the formulation alone is up to 2.8**×** faster than cuSPARSE’s generic triangular solve and 3.2**×** faster than a hand-written CUDA traversal, while halving solve memory. Relative to the strongest CPU configuration, our GPU backend is 44 faster, and relative to the single-threaded GRG library on which it builds, it is up to 473 faster. Mikado achieves order-of-magnitude end-to-end speedup and cost reduction for both analyses at biobank scale.

These gains apply from cloud instances to supercomputer nodes. Beyond accelerating established analyses, lowering the cost of this primitive also makes it feasible to run analyses that were previously computationally impractical, such as exhaustive interaction scans across many traits and exposures. Benchmarks span simulated cohorts, the 1000 Genomes Project, and the *All of Us* Research Program [1], demonstrating gains on real-world, biobank-scale data.

Mikado uses the construction toolkit from prior work [6, 9] to build its GRGs and focuses on computing over these. Our contribution is a methodology that reformulates computation on GRGs as sparse linear algebra: (i) a reverse topological ordering that makes the adjacency matrix strictly block-lower-triangular with binary reachability, recasting the genotype matrix–vector product as a sparse triangular solve suitable for massively parallel architectures such as GPUs; (ii) a pipelined wavefront schedule that decomposes the solve into overlapping blocked SpMV; and (iii) a hardware-portable CPU and GPU backend using standard vendor primitives, with the first GRG-native BOLT-LMM-inf formulation. The sparse triangular framing is a property of the representation’s structure rather than of the GRG specifically: any DAG-structured genotype encoding admits the same reverse topological ordering and block-triangular solve, so the reformulation carries over to other DAG-based encoding schemes. Its benefit, however, is not universal: the number of sequential stages equals the longest path *H* of the DAG, so the exposed parallelism is large only when *H* is small relative to the number of nodes, as guaranteed by the multi-pass GRG construction (*H* ∼ log_2_ *n*). On a DAG with a long dependency chain, the solve degenerates to the serial case, and no scheduling strategy can fully recover parallelism.

## 2 Results

The central computational primitive in statistical genetics analyses is the genotype matrix–vector product. Given the genotype matrix *G* ∈ {0, 1} ^*n*×*m*^, where *n* is the number of haploid genomes and *m* is the number of mutations, the goal is to compute *Gv* for *v* R^*m*^ or *G*^⊤^*w* for *w* R^*n*^. Converting haploid *Gv* and *G*^⊤^*w* into corresponding diploid results is trivial [12]. Conventional approaches operate on an explicit genotype matrix, whether bit-packed and dense, as in the `.bed`

format in PLINK1, or stored in a sparse representation, as in the `.pgen` format in PLINK2. In GRG-based computation, *G* is never explicitly materialized; it remains implicit in the graph’s reachability structure. In this section, we demonstrate that GRG matrix multiplication can be implemented efficiently using sparse block triangular solves (SpTRSV), that SpTRSV can be parallelized with SpMV-based block forward/backward substitution, and that even more parallelism can be achieved with a pipelined wavefront schedule.

### 2.1 From GRG Matrix Multiply to Block Sparse Matrix-Vector Multiply

Rather than formulating the product through whole-GRG traversal [6], these products can be expressed directly as sparse linear algebra primitives on the GRG adjacency matrix. Because the GRG is a directed acyclic graph (DAG), its nodes admit a reverse topological ordering, where every node precedes its ancestors. Under any such ordering, the adjacency matrix *A* is strictly lower triangular, and the genotype matrix is a submatrix of the transpose of the reachability matrix, *T* ^⊤^ = (*I* − *A*)^−⊤^. This follows from the definition of a GRG: an individual carries a mutation if and only if the corresponding mutation node can reach that individual’s sample node, so the reachability matrix encodes the genotype relation exactly.

Concretely, the genotype matrix–vector product decomposes as

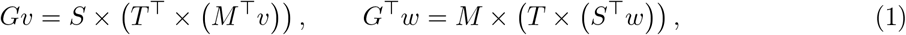

where *S* and *M* are selector matrices that map between graph nodes and samples or mutations, respectively. The selector multiplications are inexpensive scatter and gather steps; the actual work is the solve in between. The computational core applies *T* = (*I* − *A*)^−1^ or its transpose to a vector: a sparse lower-triangular solve (SpTRSV) by forward substitution for *G*^⊤^*w*, and the corresponding upper-triangular solve by backward substitution for *Gv*. For diploid analyses, the mapping between haplotype rows and per-individual genotype dosages is a linear operation absorbed into *S* and *S*^⊤^, so no extra passes over the graph are needed in either direction.

For general DAGs, the entries in *T* can be any natural number. However, because GRG is a multitree DAG [6], there is at most one path between any two nodes, so *T* is guaranteed to be binary, which is consistent with *G* being binary. In addition, since *A* is binary, we can avoid materializing the values for the nonzero elements during SpTRSV using virtual memory (Methods).

Our reverse topological ordering is based on node *height*, defined for a node in a DAG as the length of its longest path to a leaf. Ordering nodes by height gives a valid reverse topological ordering, since an edge from *u* to *v* requires *u* to have strictly greater height than *v*. Any such ordering is called a *level-set ordering*, following prior SpTRSV work [29, 30]. The ordering is not unique, since nodes of equal height may be arranged arbitrarily; that flexibility is used below to recover locality. Partitioning nodes by height gives *H* + 1 levels, where *H* is the length of the longest directed path in the GRG, that is, the maximum node height (typically *H* ∼ 30 for biobank-scale GRGs using real data). Under any level-set ordering, *A* inherits a (*H* +1) (*H* +1) block triangular structure, meaning edges run only from higher to lower levels. Because level sets in GRGs are sparsely connected, each block of *A* is itself a sparse matrix, and the solve decomposes into a sequence of sparse matrix–vector multiplications (SpMVs) and additions. By treating each block as a unit, we turn the per-edge scalar operations into per-level-pair SpMVs. The forward substitution propagates values up the graph for *G*^⊤^*w*, and backward substitution propagates them down the graph for *Gv*.

Each SpMV is internally parallel, but contributions to the same destination level execute serially, so the level-set schedule requires one synchronization stage per level. This is the level-set approach to SpTRSV [29, 30], with our height-based ordering as the analysis phase and block forward/backward substitution as the solve phase. The level-set ordering achieves the minimum possible number of sequential stages, *H*, which is equal to the length of the critical path. Thanks to the multi-pass construction of the GRG [6], *H* grows only as ∼log_2_ *n* with cohort size *n*. As a result, the block structure reduces the sequential depth from Θ(*K*), incurred by scalar forward and backward substitution that ignores block structure, to Θ(*H*) ∼ Θ(log_2_ *n*), which is below log_2_ *K*.

Other reverse topological orderings expose less parallelism; the GRGL baseline, for instance, uses a postorder DFS labeling, which is itself a valid reverse topological ordering that favors single-threaded cache locality but can leave many nodes on a serial chain, which is a poor fit for GPUs (Methods). To maximize locality in our level-set ordering without compromising parallelism, we break ties among equal-height nodes by comparing their postorder DFS labels. Empirically, this gives the blocks of *A* a sparse structure that makes SpMV efficient (Methods). Finally, writing *A*_*km*_ for the block that maps source level *m* to destination level *k*, this block can be dispatched as soon as level *m* is finalized, rather than waiting for all levels *l* ≤ *k* − 1. Mikado, our methodology and its CPU and GPU implementation, exploits this with a pipelined wavefront schedule that combines the simplicity of level-set methods [29, 30] and the parallelism of synchronization-free approaches [31]. On CPUs, it uses sparse routines from the Intel Math Kernel Library [32]; on GPUs, it uses sparse routines from cuSPARSE [27]. The block-triangular solve is described in full in Methods.

### 2.2 Kernel-Level Performance

To separate the gain of the formulation from the gain of the hardware, Mikado GPU is evaluated on the same GPU against two baselines that operate on the same graph: (i) a direct SpTRSV of the same system using cuSPARSE’s `cusparseSpSV` API, and (ii) a hand-written, graph-first CUDA implementation of GRG traversal with the load-balancing and memory-staging optimizations described in Methods. On five chromosomes of the simulated 500k cohort at *k* = 1, Mikado GPU is 3.2 faster than graph-first in both directions and 2.8 (upward) and 1.3 (downward) faster than direct SpTRSV (Fig. 2a,b). The gap over graph-first, which also partitions nodes by height, isolates the benefit of expressing each level pair as a vendor-tuned SpMV with coalesced, load-balanced access rather than as a handwritten per-node kernel; the gap over direct SpTRSV isolates the benefit of the blocked, pipelined schedule over cuSPARSE’s generic analysis-phase solver. Direct SpTRSV also requires a separate external buffer for each direction, each at least as large as the sparse matrix itself, more than doubling memory usage. The other approaches, including Mikado GPU, reuse a single stored format for both directions and, given the same additional memory, could optimize further, for instance by materializing the transposed matrix explicitly.

**Fig. 1.**
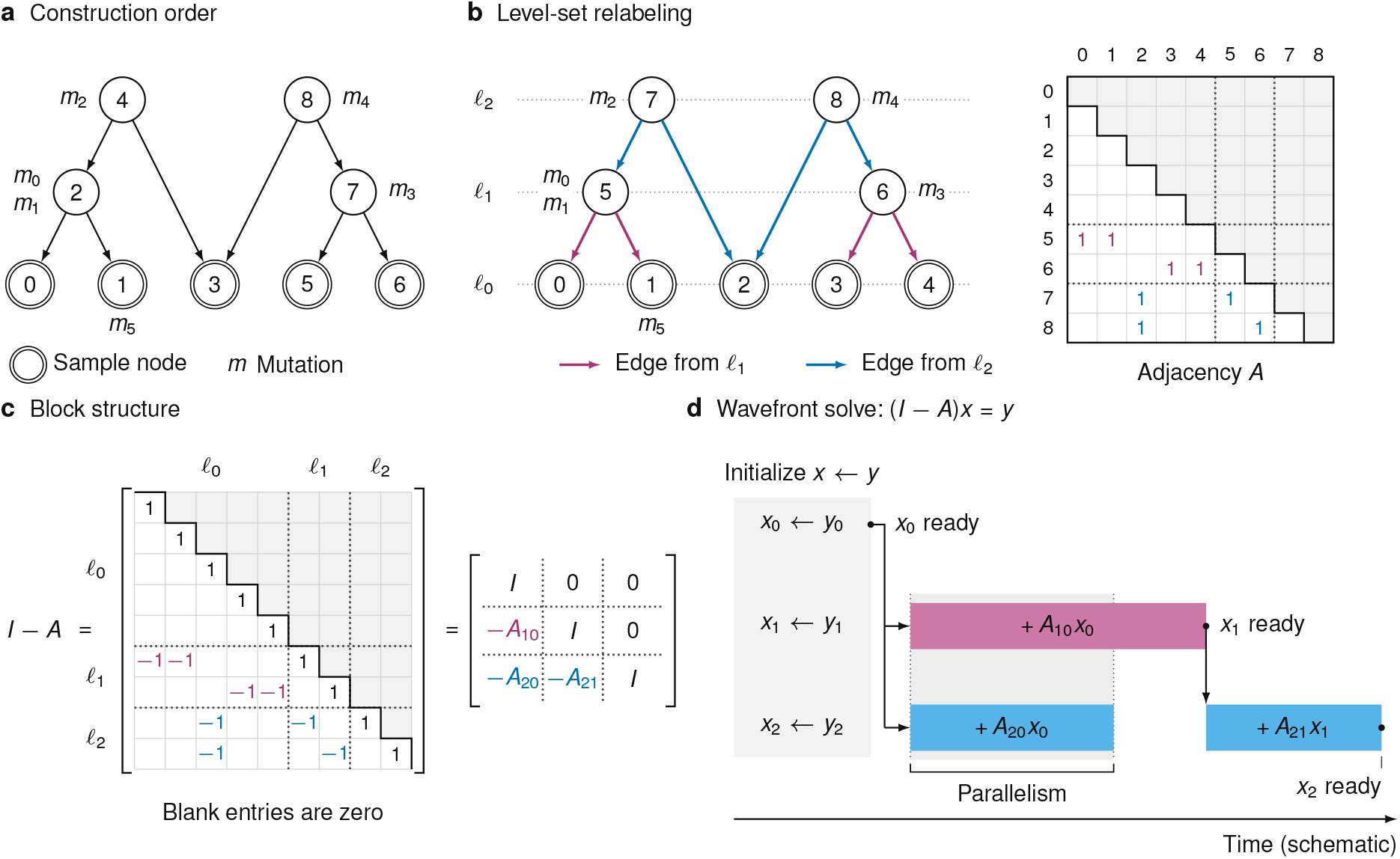
Our level-set relabeling exposes a block triangular matrix that enables efficient pipelined execution. (**a**) A toy GRG with five samples (double circles) and four ancestral nodes (single circles), carrying mutations *m*_0_–*m*_5_; edges point from ancestors to descendants. The nodes are numbered in the DFS postorder generated during graph construction, which ensures that every edge goes from a higher to a lower index. (**b**) The same graph after level-set relabeling, with nodes of equal height renumbered to have contiguous indices and dotted horizontal guides indicating the levels. The edges from ancestors in *l*_1_ are purple and those from ancestors in *l*_2_ are blue, matching the colored ones in the strictly lower-triangular adjacency matrix *A*. Dotted matrix dividers mark the level blocks. Blank cells are zero; gray marks the structurally zero region on and above the diagonal. (**c**) The triangular system matrix *I* − *A*, shown first entrywise and then in block form. Dotted dividers partition both representations by levels *l*_0_, *l*_1_, *l*_2_. Black diagonal entries form identity blocks; colored off-diagonal −1 entries form the blocks *A*_10_, *A*_20_, and *A*_21_. (**d**) A schematic wavefront solve of (*I* − *A*)*x* = *y*, with rows for *x*_0_, *x*_1_, and *x*_2_. The initialization *x* ← *y* shown across all three rows finalizes *x*_0_ and initializes the other destination vectors, *x*_1_ and *x*_2_. Colored bars indicate updates on a single ordered stream per destination level; each bar represents one SpMV. The updates using *A*_10_*x*_0_ and *A*_20_*x*_0_ overlap during the shaded time interval. The update using *A*_21_*x*_1_ begins after *x*_1_ is finalized and the preceding update to *x*_2_ has completed, yielding *x*_2_ = *y*_2_ + *A*_20_*x*_0_ + *A*_21_*x*_1_.

**Fig. 2.**
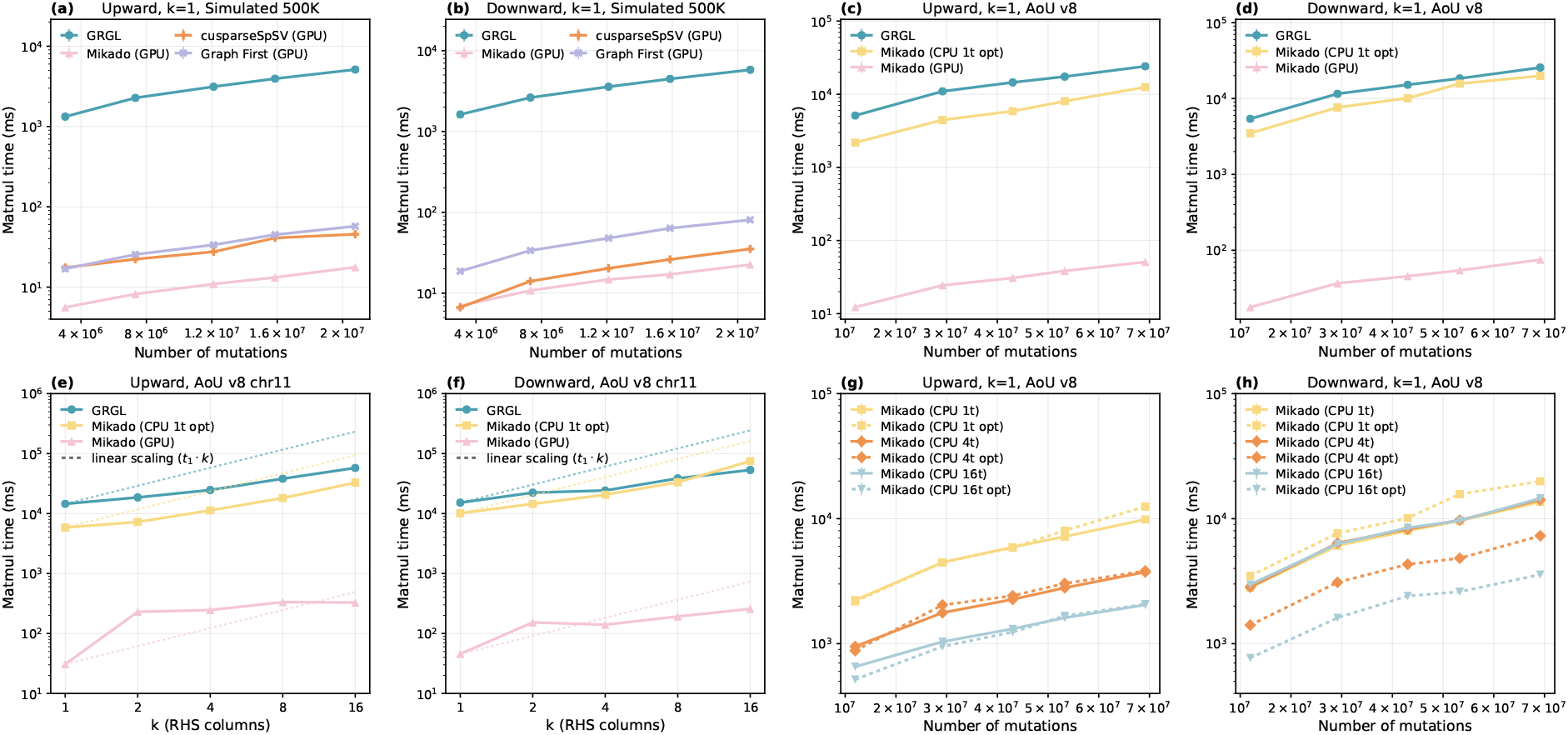
Performance of the matrix–vector primitive. Each panel pair shows upward (left) and downward (right) traversal. MKL’s `optimize` mechanism (Methods) is enabled for the Mikado CPU curves in **(c**,**d)** and **(e**,**f). (a**,**b)**: Primitive runtime on five chromosomes from the simulated 500k cohort at *k* = 1. GRGL, Mikado (GPU), cuspars-eSpSV (GPU), and Graph First (GPU) are shown; all A100 measurements in (a, b) use 40 GB devices. cusparseSpSV (GPU) uses the `SpSV` primitive provided by cuSPARSE. Graph First (GPU) is a hand-written, highly optimized, traversal-based implementation (Methods); **(c**,**d)**: Primitive runtime across five *All of Us* v8 chromosomes at *k* = 1 for both upward and downward traversal. GRGL, Mikado (CPU 1t opt), and Mikado (GPU) are shown. A100 measurements in (c–f) use 80 GB devices. **(e**,**f)**: Right-hand side (RHS) width scaling performance of GRGL and Mikado on *All of Us* v8 chromosome 11. Here, *k* is the number of RHS columns propagated per traversal. For Mikado GPU, increasing *k* from 1 to 2 multiplies runtime by 7.5 upward and 3.4 downward, whereas increasing *k* from 2 to 16 multiplies runtime by only 1.4 and 1.7, respectively. **(g**,**h)**: Mikado CPU scaling on the same *All of Us* chromosomes at *k* = 1, using 1, 4, and 16 threads, with and without optimization. Optimization requires preprocessing time and memory (Methods). For a fixed thread count, optimization yields geometric-mean downward speedups of 2.0× at four threads and 3.8× at 16 threads (3.5–4.1× across chromosomes), at the cost of a 24–63% higher single-thread runtime; upward benefits are small and variable.

For users of the existing GRG software, the impact is substantial. On an NVIDIA A100 80 GB GPU, across five *All of Us* chromosomes at *k* = 1, Mikado GPU achieves a geometric-mean speedup of 452× for upward multiplication (*G*^⊤^*w*) and 326× for downward multiplication (*Gv*) over the single-threaded CPU GRG baseline [6], with a peak upward speedup of 473× (Fig. 2c,d). GRGL is single-threaded within a chromosome, so this is the like-for-like baseline. Compared with the strongest CPU configuration tested here, the 16-thread optimized Mikado CPU backend, the GPU advantage is on average 44× (Fig. 2c,d,g,h), and the cost comparison in the next section normalizes for the hardware differences. The optimized single-threaded Mikado CPU backend achieves geometric mean speedups of 2.3× and 1.4× over GRGL in the upward and downward directions, respectively.

On *All of Us* chromosome 11, Mikado GPU is the fastest at every tested right-hand side (RHS) width (Fig. 2e,f). Going from *k* = 1 to *k* = 2 increases GPU runtime by far more than 2× (7.5**×** upward, 3.4**×** downward), because `cusparseSpMM` selects a specialized kernel for *k* = 1 and a general SpMM kernel otherwise; for *k* = 2, runtime grows sublinearly in *k*, so block multiplications amortize well. GRGL overtakes the optimized single-threaded Mikado CPU backend for downward multiplication at *k* = 16, taking 53.3 seconds compared with 74.8 seconds. Fig. 2g,h shows Mikado CPU performance under different thread and optimization configurations. Going from 1 to 16 threads without MKL optimization gives a geometric-mean upward speedup of 4.3 but essentially no downward speedup; optimization enables downward scaling but slows single-threaded downward execution.

Results on two additional machines with NVIDIA V100 and GH200 superchip, respectively, further demonstrate the efficiency and portability of Mikado‘s formulation (Extended Data Fig. 1). AMD GPU kernel results are reported in Extended Data Fig. 1, with experimental details in Supplementary Note 3. The end-to-end runtime benefits from these kernel-level gains in proportion to the fraction of total time the primitive consumes. This fraction is large for the iterative analyses below but small for a single-pass workload such as GWAS (Figs. 3, 4, and Extended Data Fig. 3).

**Fig. 3.**
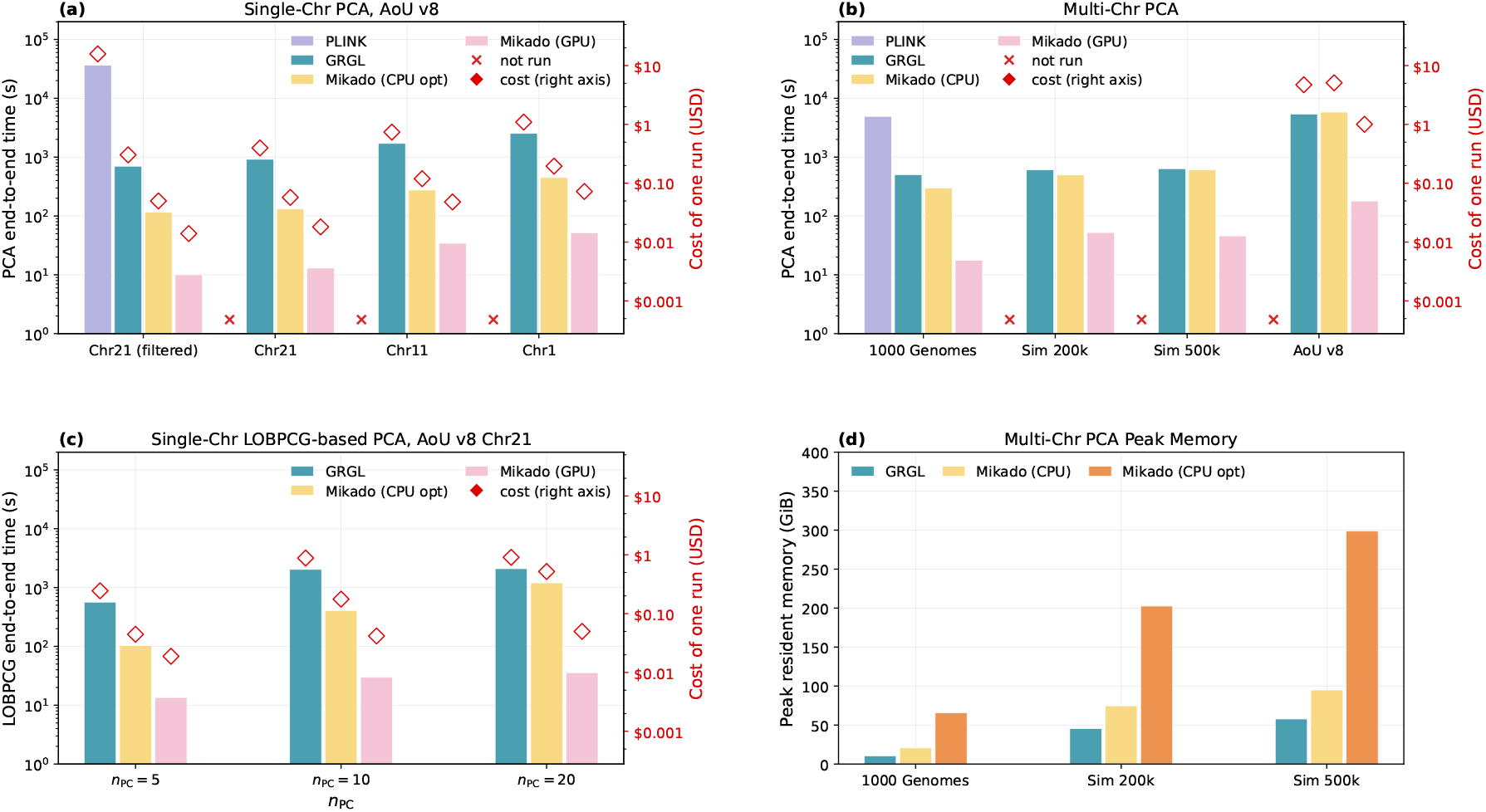
Runtime and cost of PCA. Time is shown on the left axis. The corresponding cost on the Researcher Workbench is shown on the right axis. A100 measurements in (a, c) use 80 GB devices. **(a)** A single-chromosome PCA on *All of Us* v8. `chr21(filtered)` removes non-SNP and multiallelic variants. Mikado CPU and PLINK use 16 threads. MKL’s `optimize` is enabled and optimization time is included. PLINK failed to compute results within the 72 hours time limit, except for chr21 (filtered). **(b)** Multi-chromosome PCA on four different datasets. Mikado CPU uses 1 thread per chromosome; Mikado (GPU) uses 4 GPUs. Mikado (GPU) uses A100 40 GB devices for 1000 Genomes and the simulated cohorts, and 80 GB devices for *All of Us*. MKL’s `optimize` is disabled. PLINK failed to compute results within the time limit, except for 1000 Genomes. **(c)** Single-chromosome LOBPCG-based PCA on chr21 from *All of Us* v8. Mikado CPU uses 16 threads. MKL’s `optimize` is enabled. LOBPCG begins by multiplying the genotype matrix by a block of *n*_PC_ columns; the block size decreases as individual eigenpairs converge and are deflated. **(d)** CPU memory usage for multi-chromosome PCA across GRG-based backends. GRGL uses the least memory. Mikado CPU with the MKL backend uses 1.6–2.0× as much memory, reflecting the absence of delta encoding in the `.grg_spmv` format. Mikado CPU memory usage increases to roughly 4.5–6.3× that of GRGL when MKL optimization is enabled.

**Fig. 4.**
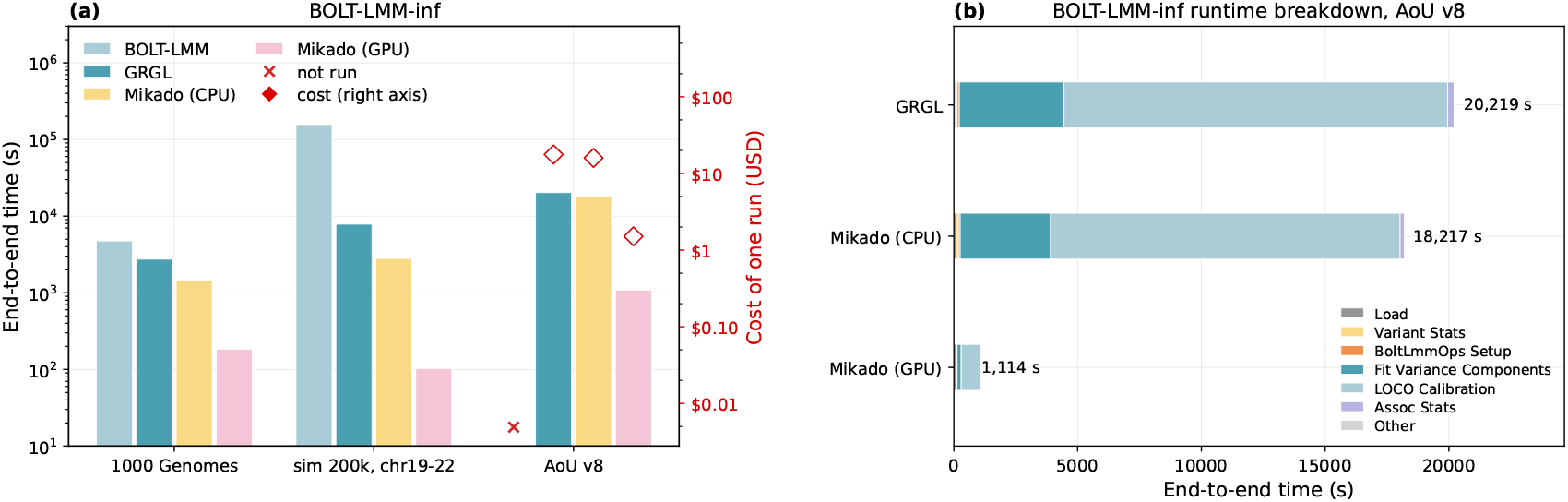
Runtime and cost of BOLT-LMM-inf. These datasets are filtered as described in Methods. **(a)** The multi-chromosome BOLT-LMM-inf runs. The three datasets are further filtered. Output write time is excluded for all implementations. Mikado CPU uses 1 thread per chromosome; Mikado (GPU) uses 1 A100 GPU. Mikado (GPU) uses 40 GB device for 1000 Genomes and the simulated cohort, and 80 GB device for *All of Us*. MKL’s `optimize` is disabled. The official BOLT-LMM is allowed 64 threads; on the *All of Us* dataset it is prohibitively expensive to run. **(b)** Detailed end-to-end runtime breakdown of BOLT-LMM-inf runs on the *All of Us* dataset. A single run is selected for each backend. Mikado (GPU) uses an A100 80 GB device.

### 2.3 Biobank-Scale Analysis Applications

Mikado accelerates biobank-scale genotype matrix–vector products by up to 473× over the single-threaded CPU GRGL baseline [6], and this primitive-level gain translates into large end-to-end speedups for the two iterative analyses presented here, PCA and BOLT-LMM-inf; the single-pass case, GWAS, is reported in Extended Data Fig. 3. The end-to-end speedup depends on the proportion of runtime spent in matrix–vector products, since non-kernel work such as I/O and preprocessing is unaffected by the accelerated primitive; the larger this proportion is, the greater the end-to-end gain.

#### 2.3.1 Principal Component Analysis (PCA)

PCA extracts orthogonal directions of maximum variance and is widely used in population genetics to characterize ancestry and control for population stratification. Randomized PCA is typically used at biobank scale, and reduces to repeated genotype matrix–vector products (Methods). On the full *All of Us* v8 phased cohort, Mikado (GPU) computes the top 10 principal components in 180 seconds at $1.00 per run, compared with 5,392 seconds at $4.72 for GRGL, making it roughly 30× faster and 4.7× cheaper (Fig. 3b). These figures include loading and setup, with staging excluded for every backend (Methods). The computation itself accounts for only $0.14 of the Mikado (GPU) cost, about 25 seconds on four A100s, and the end-to-end time is dominated by loading the 197 GB of per-chromosome GRGs (Supplementary Table 1), indicating that Mikado (GPU) shifts the bottleneck from computation to data loading, that is, parsing the GRGs from host memory and transferring them to the devices. For GRGL, computation accounts for $4.52 of the $4.72. These PCA benchmarks use every variant in the call set. PCA for stratification control is often run on an LD-pruned subset of common variants, which reduces PLINK2’s workload by orders of magnitude. LD pruning, however, requires dataset-specific parameters, and a leave-one-chromosome-out (LOCO) PCA, which avoids pruning by using all variants on the other chromosomes, has recently been proposed [9]. That approach runs a full-variant, multi-chromosome PCA for each chromosome, which is the workload measured in this work. The comparison here measures the cost of the full-variant product required by iterative methods such as BOLT-LMM, heritability estimation, and multi-trait or interaction scan. The simulated 200,000- and 500,000-individual cohorts show roughly 10× over GRGL, while PLINK2 completes PCA only on 1000 Genomes, where it is 278× slower than Mikado (GPU). Mikado with MKL offers only marginal improvement over GRGL because the multi-chromosome workload is memory-bound and the formulation does not reduce the number of memory accesses. AMD GPU results for the simulated cohorts are presented in Supplementary Table 9. For single-chromosome experiments, Mikado achieves larger speedups over GRGL, reaching up to 71× on GPU and 6.9× with MKL (Fig. 3a). GRGL does not support multithreading within a single chromosome, whereas Mikado‘s formulation exposes intra-chromosome parallelism. This is where our CPU backend adds value, since with one thread per chromosome, the two are nearly equivalent (Fig. 3b).

PLINK2 cannot complete PCA on any full chromosome within the 72-hour limit. On filtered chromosome 21, it finishes in 36,452 s, GRGL in 701 s, and Mikado (GPU) in 10.0 s, with the costs shown in Fig. 3a. The 3,650× end-to-end speedup of Mikado (GPU) over PLINK2 can be factored into two stages,

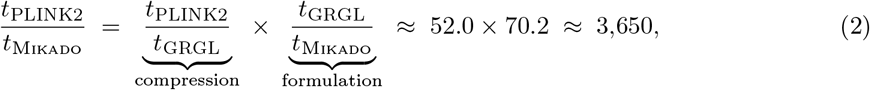

where the first factor is the GRG’s compression, which reduces both memory footprint and arithmetic work, and the second is the impact of this work: recasting GRG traversal as sparse linear algebra primitives that exploit the GPU’s parallelism and memory bandwidth^2^.

Under the block-vector LOBPCG solver, Mikado (GPU) consistently outperforms GRGL, while Mikado CPU’s advantage shrinks as the number of principal components increases, in line with the kernel benchmark (Fig. 3c). Unlike the default `eigsh` solver, which multiplies the GRG by a single vector at each step, LOBPCG multiplies it by a block of up to *n*_PC_ columns at each step. Therefore, this panel shows the backend performance for applications that benefit from block multiplication.

#### 2.3.2 BOLT-LMM Infinitesimal (BOLT-LMM-inf)

BOLT-LMM-inf [10] is the infinitesimal variant of BOLT-LMM, widely used for biobank-scale GWAS in settings where unmodeled relatedness or population structure would otherwise inflate association test statistics. Its heritability estimation and per-variant calibration stages both rely on a conjugate gradient solver that invokes the matrix–vector primitive once per iteration (Methods), making BOLT-LMM-inf the most matrix–vector-intensive of the applications in this work.

On the full *All of Us* v8 phased cohort, filtered to 208,382 individuals and 44.6 million variants (Methods), Mikado (GPU) completes a genome-wide BOLT-LMM-inf analysis of standing height under 19 minutes on a single A100, compared with 5.6 hours for GRGL on a 24-core CPU instance, an 18.7× speedup at 11.7× higher cost-effectiveness (Fig. 4a). In contrast, the official BOLT-LMM implementation using PLINK matrix multiplication is prohibitively expensive to run at this scale.

The same code evaluated on the 22 chromosomes of the 1000 Genomes Project [33] and on chromosomes 19–22 of the simulated 200,000-individual cohort, where the official BOLT-LMM can still be run, establishes both correctness and scalability. In the CUDA comparison on the simulated cohort, which is the only dataset with a synthetic ground truth, the estimated heritability is 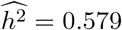 for the official BOLT-LMM and 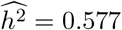 for the Mikado (GPU) implementation, compared with the ground truth value *h*^2^ = 0.6. The two estimates agree within the Monte Carlo variability of the procedure, indicating that Mikado (GPU) reproduces BOLT-LMM’s inference rather than merely its runtime, as expected from the GRG being a lossless representation. Furthermore, the *χ*^2^ test statistics from the Mikado implementation match the official BOLT-LMM v2.5 values up to a global multiplicative constant, which is attributable to differences in the pseudorandom number generator (PRNG) implementation used for Monte Carlo trace estimation and calibration-SNP selection (Extended Data Fig. 4, Supplementary Note 4.6).

For 1000 Genomes, Mikado (GPU) finishes in 183.3 s, GRGL in 2,737.4 s, and the official BOLT-LMM in 4,742.0 s (Fig. 4a). On the four-chromosome simulated 200,000-individual cohort, the three runtimes are 102.9 s, 7,840.0 s, and 152,283.4 s. The end-to-end speedup over BOLT-LMM factorizes as in Eq. (2) into the GRG’s compression and Mikado‘s formulation:

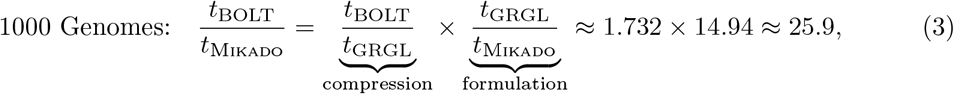

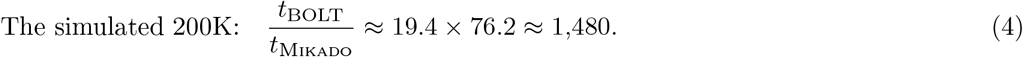

Both factors grow with cohort size: compression improves from 1.7× to 19.4× because GRG size increases sublinearly with the number of individuals, and the formulation’s share increases from 14.9× to 76.2×. The speedup over the official implementation therefore increases with the cohort size. On *All of Us*, Mikado CPU achieves a 1.1× speedup over GRGL on the same hardware, yielding 1.1× more work per dollar, whereas Mikado (GPU) on an A100 instance is 18.7× faster at 1.6× the hourly cost, resulting in the 11.7× cost-effectiveness gain noted above.

## 3 Discussion

Recasting GRG traversal as sparse linear algebra changes what biobank-scale population genetics can achieve and at what cost, because a wide range of analyses reduce to the single primitive it accelerates: genotype matrix multiplication. Randomized and truncated PCA reduce to repeated genotype matrix–vector multiplies against the standardized genotype matrix [34, 35]; mixed-model association solvers such as BOLT-LMM invoke the same primitive within their conjugate-gradient iterations [10]; and model-based ancestry estimators, including ADMIXTURE [36, 37], alternate updates dominated by products of the genotype matrix, or its transpose, with the parameter vector. Even sparse nonnegative matrix factorization methods like sNMF [38], which rely on matrix–matrix multiplication, can benefit from iterating over the columns of the right-hand-side matrix, since Mikado reduces the arithmetic of such compute-bound products by sharing intermediate values across graph nodes. Thus, any method built on this primitive inherits Mikado‘s throughput and cost advantages by substituting the GRG-backed version for its dense or PLINK2-based counterpart, without changing the surrounding statistical model.

Methodologically, expressing the computation as a standard primitive lets the approach inherit performance and portability from a rapidly evolving accelerator software stack, such as cuS-PARSE [27], without requiring ongoing custom kernel engineering. This matters because the computing and accelerator ecosystem is now changing rapidly [39], and hand-maintained code is a recurring liability and source of effort, not a one-time cost. Relying only on standard sparse BLAS, the same Mikado code runs unchanged on a metered cloud configuration (*All of Us* Researcher Work-bench on Google Cloud, GPU and CPU instances), on the Perlmutter supercomputer at NERSC, and across four NVIDIA GPU generations, from the V100 to the tightly coupled GH200 superchip (Extended Data Figs. 1 and 2). Portability to AMD GPUs is demonstrated in Extended Data Fig. 1 and in Supplementary Note 3. Because Mikado is expressed in standard primitives rather than a bespoke kernel, the mature sparse triangular solve (SpTRSV) literature [40–42] is directly applicable, making the adoption of a faster scheduler a drop-in improvement rather than a redesign.

On a single GPU, the core product accelerates by up to 473× over the single-threaded CPU GRGL baseline [9], but the end-to-end speedup is governed by the fraction of the runtime spent on matrix–vector work. Computations with few matrix–vector multiplies, such as a single GWAS pass, are dominated by chromosome loading and improve only marginally, whereas PCA and BOLT-LMM-inf place the multiply in a batch and/or an iterative inner loop and achieve order-of-magnitude end-to-end speedups. Operating on cloud infrastructures required for biobank data, such as *All of Us*, makes cost a first-class concern, not just runtime, because the same budget then enables more analyses. Because Mikado finishes much sooner, the shorter runtime offsets any hourly-rate premium, so accelerated analysis is typically both faster and cheaper, although the relative ratios vary by workload. The ratios of time and cost saved, rather than the absolute values, are what carry over to other scenarios: a study spanning thousands of phenotypes, G E exposures, or LOCO segments inherits these savings, multiplied across every run. Big data and machine learning research already benefit in speed and cost from modern accelerators; Mikado extends these advantages to population genetics, where irregularity and sparsity have kept them out of reach.

The reformulation is not specific to the GRG. Any DAG-structured genotype representation admits the same sparse triangular framing, so the performance and portability advantages carry over.

The linear ARG [43] is a recent, closely related encoding whose computation also reduces to a sparse triangular system, so our two-step reformulation applies to it as well. Mikado is designed for GRGs, whose multi-tree construction keeps graph height small. The speedup on other encodings depends on their structure, because the height determines the number of sequential stages in the solve and therefore the parallelism that Mikado‘s level-set blocking and pipelined wavefront schedule can expose. Because the linear ARG is a general DAG rather than a multi-tree, diamond-shaped paths require signed edge weights, so it requires a general, rather than binary, SpTRSV. For ancestral recombination graphs (ARGs), tskit’s genetic relatedness vector products [44] leverage the tree sequence encoding to efficiently compute an implicit linear operator. ARG-RHE [45] shows that ARG topology enables sublinear-time genotype matrix–vector products on CPUs, demonstrating that shared genealogical structure provides asymptotic gains over tabular formats. The present contribution is complementary, mapping analogous GRG structure to sparse primitives suited to GPU acceleration and delivering large constant-factor speedups at biobank scale. More broadly, efforts such as traceax [46], a JAX-based [47] framework for stochastic trace estimation, accelerate matrix–vector multiplication on explicit genotype matrices. In contrast, the approach here implicitly accelerates the product on the compressed GRG, so any downstream estimator, whether trace-based or not, inherits the speedup.

The main limitation of Mikado is the device memory ceiling: single-node analysis is restricted to cohorts whose per-chromosome GRGs fit within the total available GPU memory. Under our quality control pipeline, the largest per-chromosome `.grg_spmv` file for the full *All of Us* cohort (414,830 individuals) is 16 GB, so any single chromosome fits on a 24 GB GPU, and the full 197 GB cohort requires four 80 GB A100s to be resident simultaneously (Supplementary Table 1). The CUDA host-memory streaming mode that transfers blocks on demand (Methods, Extended Data Fig. 2), as well as partitioning across nodes for larger cohort sizes, are possible routes past this ceiling. A secondary cost is memory footprint, because the block format roughly doubles the stored size relative to GRGL’s delta encoding, which compresses repeated edges (Methods). This overhead, however, is removable rather than intrinsic, since delta encoding can also be applied to the block format; exploring this direction is left for future work.

The scientific impact of removing this compute bottleneck is that some analyses become feasible, not just faster. Gene–environment and gene–gene interaction scans are a clear example, because an exhaustive scan multiplies the number of genotype matrix products by the number of exposures and candidate loci; current studies therefore restrict themselves to a handful of exposures and traits [48] or to a preselected set of loci [49, 50]. Reducing the cost per product by an order of magnitude lets the same budget to cover an order of magnitude more exposures, traits, or loci, bringing exhaustive G×E scans across many exposures and traits, as well as three-way G×G×E or G×G×G scans over candidate sets, within reach at biobank scale.

### Algorithm 1

Level-set Block Forward Substitution

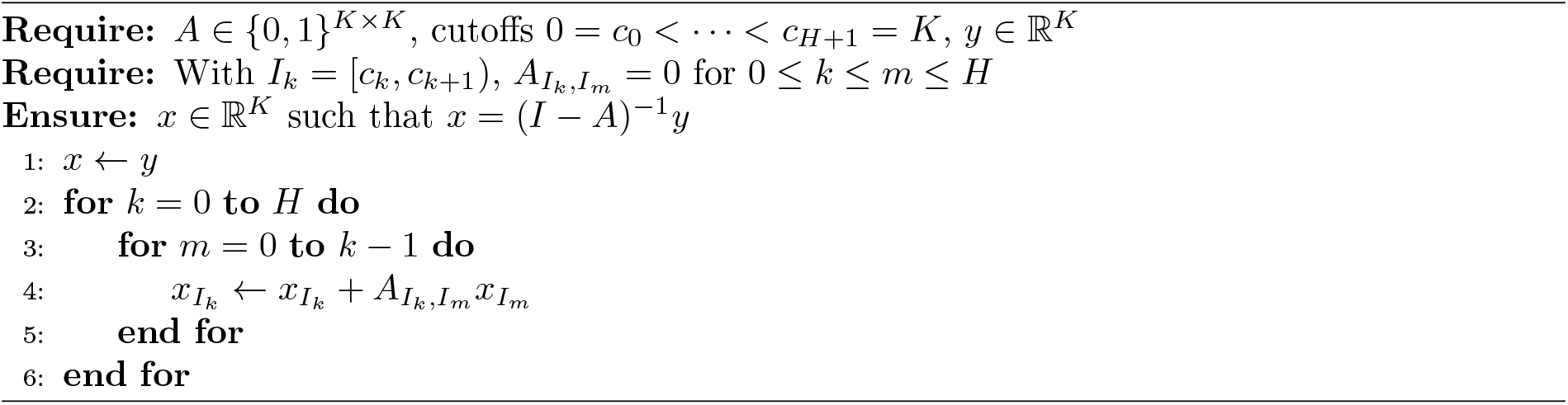

## 4 Methods

### 4.1 Genotype Representation Graph

A GRG encodes genotypes as a directed acyclic graph with a multi-tree structure [6, 9]. A multitree is a directed acyclic graph (DAG) in which a node may have multiple parents, but at most one directed path connects any pair of nodes. The samples are mapped to leaf nodes, while variants may be mapped to any node in the graph. Each variant allele is associated with a node, and paths connect samples to the variants they carry. Sample *s*_*i*_ carries variant *v*_*j*_ if and only if the node associated with *s*_*i*_ is reachable from the node associated with *v*_*j*_. For example, in Fig. 1a, the sample at node 1 carries exactly mutations *m*_0_ and *m*_1_ (from node 2), *m*_2_ (from node 4), and *m*_5_ (from node 1 itself); it does not carry *m*_4_ because no path leads from node 8, which carries *m*_4_, to node 1. This structure allows a GRG to compress genotype datasets losslessly, and GRGs can be constructed efficiently from explicit, dense genotype matrices [6, 9]. A GRG is constructed from haploid or phased genomes but supports both haploid and diploid computation.

Beyond compression, the GRG enables fast computation directly on the genotype matrix without first decompressing it. That is, multiplying the genotype matrix by a vector is equivalent to a downward traversal of the GRG, where the input vector supplies initial values at the mutation nodes. Conversely, multiplying by the transposed genotype matrix corresponds to an upward traversal, where the leaf nodes are assigned the initial values. The compression reduces both memory footprint and computation, because shared subgraphs are visited once rather than once per sample.

### 4.2 Node Ordering and Wavefront Scheduling

Our SpTRSV formulation works for any reverse topological ordering, but the specific choice trades off parallelism against locality. The baseline GRGL toolkit uses postorder DFS labeling, which is a valid reverse topological order. It keeps dependent nodes close for cache reuse but leaves many blocks with only a single node, which limits parallelism. Equation 5 highlights the block structure that partitions *A* into maximal contiguous dependency-free batches: postorder DFS yields five blocks and four sequential steps (left), whereas any level-set ordering yields only three blocks and two sequential steps (right). The former favors temporal locality; the latter favors parallelism, at the cost of cache thrashing on a machine whose cache can hold only one node at a time.

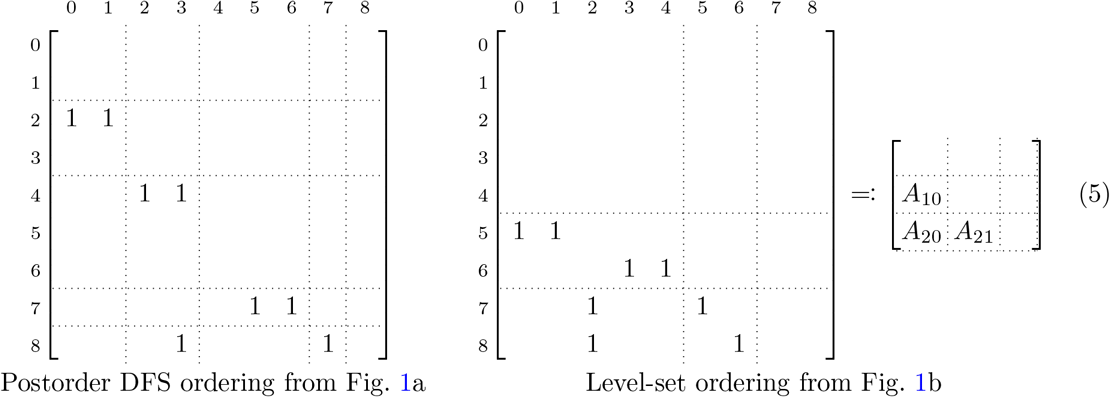

#### Algorithm 2

Pipelined Wavefront Block Forward Substitution (Mikado).

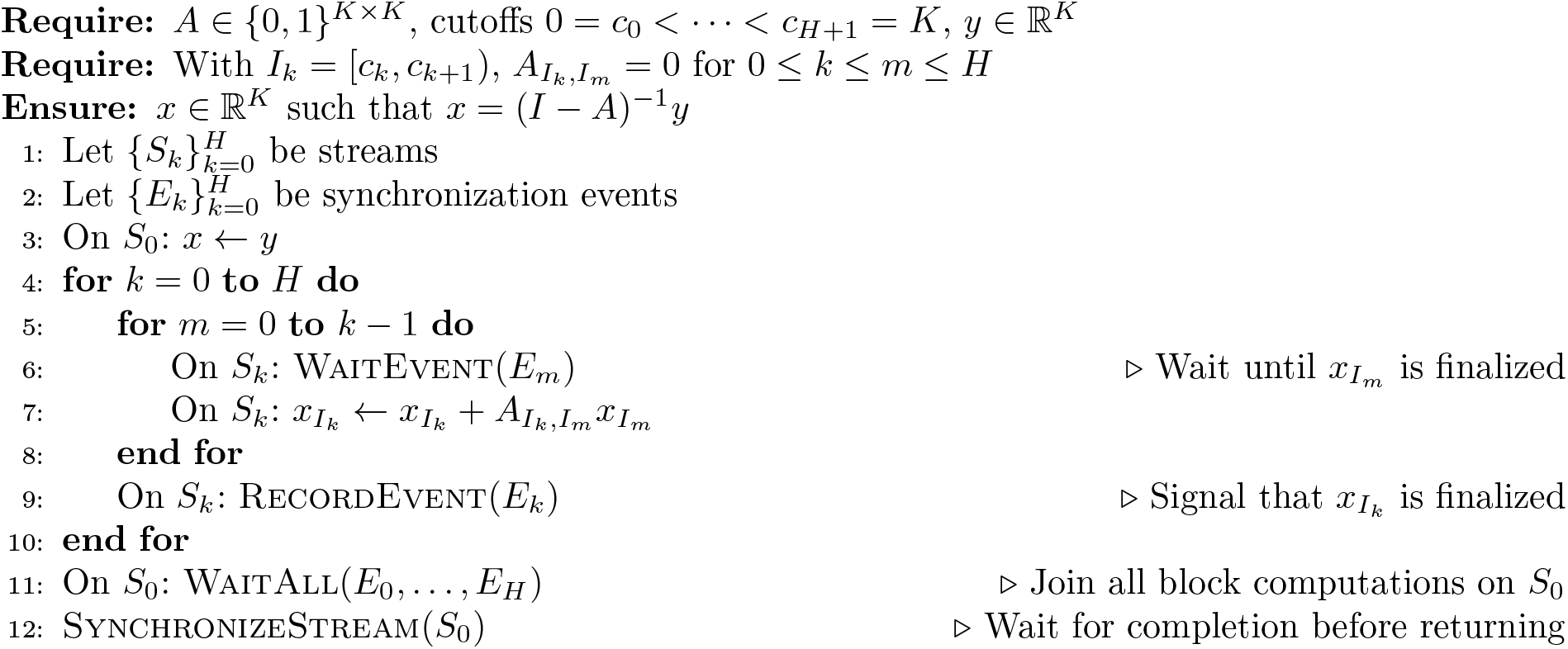

Level-set ordering minimizes the number of sequential steps, making it well suited to massively parallel accelerators like GPUs. To approach the parallelism–locality Pareto frontier, we recover locality without increasing the sequential depth by exploiting the freedom in within-level ordering: ties among equal-height nodes are broken using their postorder DFS labeling. In practice, constructed `.grg` files already carry postorder DFS labels, so Mikado only needs to stably sort nodes by height. This within-level ordering keeps each node’s dependencies close. In Fig. 1b, node 5 depends on nearby nodes 0 and 1 rather than on distant ones. The dependencies of adjacent nodes also stay close to one another: if Fig. 1b were a sub-DAG of a larger graph with many height-zero nodes, nodes 5 and 6 might be labeled 1005 and 1006, but their dependencies (0 and 1, and 3 and 4) would still be adjacent. Empirically, some blocks of *A* exhibit a nonzero diagonal pattern, improving cache locality during SpMV.

The chosen ordering reduces the genotype matrix–vector product to the block-triangular system (*I* − *A*)*x* = *y*, where *y* holds the scattered input and *x* the propagated node values. For the three-level example in Fig. 1c, this is:

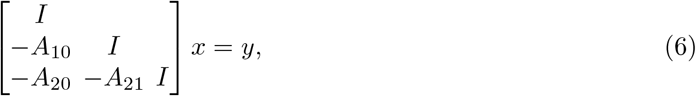

solved by block forward substitution: *x*_0_ ← *y*_0_, then *x*_1_ ← *y*_1_ + *A*_10_*x*_0_, then *x*_2_ ← *y*_2_ + *A*_20_*x*_0_ + *A*_21_*x*_1_. A level-set schedule places a global barrier between these steps, so level *k* begins only after every node in levels 0, … , *k* − 1 has been finalized (Algorithm 1). Our pipelined wavefront schedule (Algorithm 2) replaces the global barrier with source completion events and uses one stream *S*_*k*_ per destination level. The loops execute on the host, and “On *S*_*k*_“ denotes enqueuing an operation in the ordered stream *S*_*k*_. A product begins after its source is finalized and preceding writes on *S*_*k*_ have finished. Thus, *A*_20_*x*_0_ can overlap *A*_10_*x*_0_, while *A*_20_*x*_0_ and *A*_21_*x*_1_ accumulate sequentially into *x*_2_ on *S*_2_. Fig. 1d illustrates the intended overlap.

The parallelism-locality tension, and the DAG scheduling problem more broadly, are well studied [51, 52]. Because Mikado recasts GRG traversal as standard sparse BLAS, the mature SpTRSV literature applies directly. For example, existing schedulers such as SpMP [40], DAGP [41, 42, 53], and HDagg [42], along with recent advances such as GrowLocal [54], DFA-SpTRSV [55], and ElasticDivide [56], are drop-in improvements rather than a complete redesign.

### 4.3 Mikado’s Implementation

For Mikado CPU, Algorithm 1 was implemented using the inspector-executor API for sparse BLAS routines in Intel MKL [32]. The vendor SpMV routines require a matrix handle in a standard sparse format (CSC, CSR, or COO). Following GRGL’s convention, we store *A* in CSR format, or equivalently, *A*^⊤^ in CSC format. Since all nonzero entries of our blocks 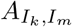 equal one, the `val` array is redundant. This redundancy was exploited by allocating a shared all-ones array, aliasing it to the `val` pointer of every block, and `mmap`ing it to span arbitrarily many logical bytes while occupying few physical bytes, greatly reducing the handle’s memory footprint. MKL’s optional `optimize` API analyzes matrix structure and applies hint-based optimizations [32], but it allocates an internal buffer roughly the size of the original matrix and is incompatible with the aliasing scheme, at least doubling memory usage; it is therefore exposed as a user option. MKL’s multithreaded single-matrix SpMV scaled unpredictably, especially with optimization disabled, so chromosome-level parallelism was prioritized following GRGL, with multiple chromosomes processed concurrently across threads. Hybrid parallelism is supported, and allocating more threads to larger chromosomes helps mitigate the straggler effect caused by load imbalance in some cases.

On GPU, two strategies implement GRG-based matrix–vector products: a graph-first baseline that ports the CPU traversal logic to the GPU, and the Mikado GPU approach, the primary contribution of this work. Both use CUDA streams on NVIDIA GPUs, share the matrix-vector primitive used by the applications, and inherit GRG construction unchanged from the original toolkit [6]. The graph-first baseline mirrors CPU GRG traversal: nodes were partitioned by height to break dependencies, and a height-level kernel propagated values between adjacent levels using warp-level reductions for bottom-up traversal and `atomicAdd` for top-down traversal. It was optimized in three ways:

1. To amortize per-block scheduling overhead, each thread block processed multiple same-level nodes when *k* = 1. For larger *k*, each thread block handled a single node, with consecutive threads assigned to the same down edge for coalesced access;
2. For *k* ≠ 1, a node’s edges were staged in a shared memory buffer before accessing their values, overlapping edge loading with computation;
3. Threads per node were assigned dynamically to balance the load within each height partition, with a dedicated outlier kernel giving high-degree nodes extra threads.

This baseline depends heavily on CUDA and cannot be retargeted directly, but it is already substantially faster than the CPU implementation [6] (Fig. 2a,b).

The Mikado GPU approach instead expresses the computation as a sequence of standard sparse primitives. The adjacency matrix *A* is stored once in a reverse topological (level-set) ordering that makes it strictly lower triangular, partitioned into height-based blocks 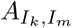 , and Algorithm 2 was implemented by dispatching off-diagonal blocks to `cusparseSpMM` routines running in concurrent CUDA streams, orchestrated with the CUDA events API. By contrast, cuSPARSE’s SpTRSV API (`cusparseSpSV`, the direct SpTRSV baseline in Fig. 2a,b) requires a costly analysis phase and an external buffer empirically at least as large as the matrix, both repeated for each solve direction. Our block forward and backward substitution, however, precompute the level structure once and reuse a single stored *A* for both directions, eliminating per-direction preprocessing and cutting the required solve memory in half. The virtual memory aliasing described above was applied on the device using the CUDA Virtual Memory Management API. On CUDA, for datasets whose GRGs exceed a single GPU’s memory, whole per-chromosome GRGs were assigned to separate devices via a device map rather than partitioning a single GRG. NVLink is used when available, since GPU-to-GPU communication can otherwise dominate runtime. For GRGs too large to fit in device memory, a CUDA streaming mode transfers blocks on demand from host memory using double buffering, so one block is processed while the next is in transit (Extended Data Fig. 2). Because Mikado relies only on standard sparse BLAS, it can be retargeted across hardware without vendor-specific rewrites, as demonstrated by our ROCm support for AMD GPUs (Supplementary Note 3), whereas the graph-first baseline, written in CUDA, cannot be retargeted without substantial reengineering.

The block formulation reorganizes the nodes and edges of the native GRG format into a block format using a lightweight conversion that requires no significant computation. Because the converted `.grg_spmv` format does not yet support delta encoding, its files are larger than the corresponding `.grg` files: on *All of Us*, `.grg_spmv` totaled 197 GB (1.9× the `.grg` size) yet was still 15× smaller than the corresponding `.vcf.gz` files.

### 4.4 Per-Application Matrix-Vector Workload

The three applications differ in how heavily they use the matrix-vector primitive. GWAS with a quantitative phenotype requires only a fixed, small number of products per phenotype (a single *G*^⊤^*w* for the per-variant score, plus one for allele frequencies shared across phenotypes), so its runtime is dominated by chromosome loading rather than by the kernel (Extended Data Fig. 3). PCA has two equivalent formulations: (i) an eigendecomposition of the genetic relationship matrix and (ii) a singular value decomposition of the genotype matrix. Critically, at biobank scale, exact eigendecom-position is infeasible, so randomized SVD or implicitly restarted Lanczos methods are used; both reduce PCA to repeated genotype matrix–vector products. The top 10 principal components of *All of Us* were computed with `grapp` using a SciPy/CuPy implicitly restarted Lanczos solver [9], and were compared against GRGL and PLINK2 `pca –approx`, which use a similar randomized block Lanczos method [2].

BOLT-LMM-inf [10] has two main steps: a restricted maximum likelihood (REML) subroutine for heritability estimation (step 1a) and per-variant calibration using leave-one-chromosome-out (LOCO; step 1b). Both rely on a conjugate gradient (CG) solver in sample space, *GG*^⊤^*x* = *y* with *x, y* ∈ ℝ^*n*^, invoking the matrix-vector primitive once per CG iteration. The total count is *O*(*k*_a_*ET* +*k*_b_(*F* +*R*)), where *k*_a_ is the average number of CG iterations during step 1a, *E* is the number of variance-parameter evaluations, *T* is the number of Monte Carlo trials, *k*_b_ is the average number of CG iterations during step 1b, *F* is the number of LOCO folds, and *R* is the number of calibration SNPs. The infinitesimal model was chosen over the more complex Gaussian mixture BOLT-LMM because (i) its *χ*^2^ statistics are essentially identical to those of GCTA-LOCO at the SNP level for polygenic traits, (ii) it is the statistically correct model for highly polygenic traits such as height, and (iii) full BOLT-LMM begins by running BOLT-LMM-inf, which accounts for ∼ 40% of total runtime [10].

### 4.5 Experimental Setup

#### Computing Hardware

Our experiments ran primarily on the *All of Us* Researcher Workbench, hosted on Google Cloud. GPU experiments used A2 Ultra instances with NVIDIA A100 80 GiB SXM GPUs, with up to four per node connected via third-generation NVLink. For single-chromosome CPU experiments, we used `n2-standard-32` instances (32 vCPUs, 16 physical cores, 128 GB); smaller instances are more susceptible to noisy neighbor interference from co-tenant workloads. For multi-chromosome experiments, we used `n2-highmem-48` instances (48 vCPUs, 24 physical cores, 384 GB), with one physical core per chromosome and enough memory to hold the chromosomes concurrently. On-demand hourly rates are listed in Supplementary Table 15. Because these rates are metered, we report the dollar cost alongside the wall-clock time, as the two do not necessarily scale together. Cost is calculated by converting runtime from seconds to hours and multiplying by the hourly rate. The two logarithmic axes have equal decade spacing, with cost limits equal to time limits divided by 3600, so USD 1 corresponds to 3600 seconds (the USD 1/hour reference). A GPU instance has a higher hourly rate than a CPU instance, but because it finishes much sooner, it is often cheaper overall. As a result, Mikado on GPU is often both faster and cheaper than the CPU GRG baseline. On identical CPU hardware with a fixed hourly rate, cost savings equal runtime savings. However, GRGL uses less memory than Mikado CPU, so it might be possible to switch to lower-tier machines and reduce costs. This depends on multiple factors, including memory bandwidth, which is typically neither specified nor guaranteed in cloud infrastructures. The Researcher Workbench does not allow local disk with controlled-tier data, so datasets reside in Cloud Storage buckets with lower bandwidth and higher network variability. To mitigate this, inputs were staged on a RAM disk for all methods when memory permitted, and staging time was excluded from the reported runtimes. The Perlmutter supercomputer was used for the 1000 Genomes [33] and simulated data. Each GPU node has one AMD EPYC 7763 (Milan) 64-core CPU, 256 GB of DDR4 memory, and four A100 GPUs connected via third-generation NVLink, and was used for both CPU and GPU runs. In addition, the kernel runtimes are evaluated on DeltaAI with Grace Hopper GH200 superchips and on SDSC Expanse V100 nodes. Each GH200 superchip has a 72-core Grace ARM CPU and an NVIDIA H100 GPU, connected by NVLink-C2C (900 GB/s bidirectional). Each SDSC Expanse V100 node has two Intel Xeon Gold 6248 CPUs, and four NVIDIA V100 SXM2 GPUs. The memory capacities of both platforms are listed in Supplementary Table 15. Every single-chromosome experiment uses one GPU; multi-chromosome experiments use either one or four GPUs, as noted for each.

#### Computing Software

Mikado uses Intel MKL on CPUs, cuSPARSE on NVIDIA GPUs, and rocSPARSE on AMD GPUs. For experiments run on NVIDIA GPUs, we use CUDA 13.3.1 with cuSPARSE 12.8.2.51. This version or newer is required, as earlier releases contain a bug we identified that affects numerical precision. Because the *All of Us* Researcher Workbench provides only CUDA 12.8, we rely on CUDA forward compatibility to load newer libraries. For the CPU, we use Intel MKL 2026.1.

For PLINK, we use the PLINK2 alpha 6.9 LM 64-bit Intel build preinstalled on the *All of Us* Researcher Workbench, and PLINK2 alpha 7.1 64-bit AVX2 Intel on all other platforms. For BOLT-LMM, we built version 2.5 from source, linked against Intel MKL. GRGL 2.9 and a slightly modified version of `grapp` were used^3^. The AMD portability experiments, including their environment, workload configuration, and reproducibility information, are described in Supplementary Note 3.

### 4.6 Dataset, Quality Control, and Statistical Analysis

The primary dataset was the srWGS statistical phasing call set in the *All of Us* CDRv8 release, as phased data are recommended for GRG construction [6]: 414,830 individuals and 870,337,413 variants across 22 chromosomes. GRGs were constructed with GRG Library v2.9 [9] using default parameters^4^ Three supplementary datasets supported smaller-scale and controlled-scaling experiments. The 1000 Genomes dataset [33] is a smaller WGS cohort consisting of 3,202 individuals and 70,692,001 variants. For controlled scaling, two simulated cohorts of 200,000 and 500,000 individuals spanned all 22 autosomes at full GRCh38 length. These cohorts were generated under the four-population `OutOfAfrica_4J17` demographic model with `stdpopsim` [57] and the `msprime` backend, drawing equal numbers of diploid individuals from CEU, CHB, JPT, and YRI.

Kernel benchmarks and GWAS analyses were run on unfiltered datasets. PCA was run unfiltered, except for filtered chromosome 21 from *All of Us* v8, where non-SNP and multiallelic variants were removed. The BOLT-LMM analyses used additional filtering applied per dataset. On *All of Us* v8, variants were filtered by minor allele frequency (MAF ≥ 5 × 10^−5^) and Hardy–Weinberg equilibrium (*p* ≥ 10^−12^), with non-SNP and multiallelic sites removed; individuals were restricted to the EUR group in the *All of Us* ancestry file, excluding those with quality-control flags, leaving 220,760 individuals and 44,626,943 variants, of which the 208,382 individuals with a height measurement were included in the BOLT-LMM analysis. For the simulated 200K cohort (last four chromosomes), variants were filtered by MAF ≥ 10^−4^ and multiallelic sites removed. For 1000 Genomes, variants were filtered by MAF ≥ 10^−2^, and non-SNP and multiallelic sites were removed.

Both GWAS and BOLT-LMM require inputs beyond the genotype data. For GWAS, phenotypes were simulated at random. For BOLT-LMM on the simulated 200K cohort, phenotypes were generated with `grg-pheno-sim` v1.5 at heritability 0.6. On *All of Us*, we used standing height (concept ID `903133`), including measured and self-reported values, with sex, age, and the top 10 principal components of the filtered subset as covariates.

The GRG-based and reference implementations were compared at two levels. At the kernel level, the genotype matrix-vector products computed by Mikado were checked element-wise against the reference GRG traversal, yielding a maximum absolute error below 10^−5^. GWAS per-variant test statistics and *p*-values were compared with PLINK2 on 1000 Genomes across all variants, yielding *R*^2^ *>* 0.999 for every statistic. For PCA, the top 10 principal components were compared against PLINK2 and against `grapp` with the GRGL backend, using per-component cosine similarity with signs aligned to resolve eigenvector sign ambiguity; Mikado attains a cosine similarity of 1.000 with GRGL for all 10 components. PLINK2 produces unstable results for PC7–PC10 across runs on *All of Us*, so the PLINK2 comparison is restricted to the top six components, for which Mikado achieves a cosine similarity of 1.000. For BOLT-LMM-inf, per-SNP *χ*^2^ statistics were compared against the official BOLT-LMM output using the squared Pearson correlation. On both the 1000 Genomes dataset and the simulated 200K cohort (last four chromosomes), the GRG-based approach attains *R*^2^ *>* 0.999.

#### Statistics and Reproducibility

Reported application timings are wall-clock times under the staging convention described above. The end-to-end times include loading and setup, while compute-only timings exclude them. The run-to-run variability on the Researcher Workbench is confounded by co-tenant interference [58, 59], which the instance choice in **Computing Hardware** partially mitigates. GRG-based computations used float64 arithmetic.

The timings in Fig. 2 and Extended Data Fig. 1 are arithmetic means of kernel calls measured within a single process, after three untimed warmup calls. Each configuration uses ten timed calls, except the *All of Us* MKL configurations and *All of Us* GRGL at *k* = 2, 8, 16, which use three. CPU and NVIDIA configurations have one recorded process each; the AMD configurations, including the same-host GRGL baseline, have three measured processes after one excluded warmup process, reported as means across processes with minimum–maximum error bars. Process counts, timed call counts, and solver iteration counts are distinct.

Repetition counts for applications varied by configuration. Each Mikado GPU (CUDA) experiment was run three times to capture GPU performance variation, with one additional breakdown run for the *All of Us* BOLT-LMM experiment. GRGL and Mikado CPU experiments were mostly run once because of their longer runtime and higher cost. The repeated CPU experiments, including multi-chromosome PCA on the simulated cohorts, showed little relative run-to-run variance, as expected for longer runs. The relative standard deviation is below 2%. AMD PCA used three measured processes per cohort, and AMD BOLT-LMM-inf used one; one warmup process per application configuration was excluded. In Supplementary Table 9, we report arithmetic means and the measured end-to-end ranges.

GRG-based PCA used NumPy’s random-number generator with seed `0` for initialization; PLINK2 PCA used its default seed. Ordinary `eigsh` PCA requested the ten largest-magnitude eigenpairs of the implicit standardized *XX*^*T*^ operator. CPU SciPy/ARPACK used `tol=0`, that is, machine-precision relative accuracy for eigenvalues, with ncv = 21 and the default maxiter = 10*n*, where *n* is the operator dimension. CuPy instead compares the Euclidean norm of its Ritz-residual estimate with a supplied absolute tolerance, using ncv = 42 and the same 10*n* iteration cap. AMD PCA used absolute tolerances of 5.1747187118964926 × 10^−5^ for 200K and 1.3151467666098284 × 10^−4^ for 500K. These values were taken from the corresponding CPU records as float64 machine epsilon multiplied by the leading eigenvalue; this recorded quantity is not ARPACK’s exact stopping test. Fig. 3c instead used LOBPCG with 5, 10 or 20 PCs, solver-specific default tolerances, and a 100-iteration cap.

Post-solve PCA verification is shared across backends, performed outside the timed interval, and distinct from the solver stopping tests. For each eigenpair (*λ, v*) of *A* = *XX*^*T*^ , the checker requires finite eigenpairs and residuals, and applies

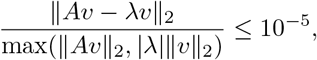

using the same threshold on the unnormalized residual if the denominator is zero. The criterion is identical for CPU, CUDA, and ROCm.

The official and GRG-based BOLT-LMM runs used seed `2026`, *R* = 30 calibration SNPs, and the automatic Monte Carlo count *T* = max(3, min(15, ⌊4 × 10^9^*/N* ^2^⌋)), where *N* is the number of analyzed individuals. The conjugate-gradient stopping criterion is the maximum, across right-hand sides, of ∥*r*∥_2_*/*∥*r*_initial_∥_2_ ≤ 5 × 10^−4^, relaxed to 5 × 10^−3^ during variance fitting. These tolerances match the official BOLT-LMM defaults; GRG-based runs used a 10,000-iteration cap, whereas the historical official runs used the 500-iteration default. The GRG-based variance fitting procedure permits up to five secant updates. If the search ends without convergence, it issues a warning and returns the best finite variance estimate retained; sampling and calibration differences are described in Supplementary Note 4.6.

#### Disclosure of AI Use

The authors take full responsibility for the study’s methods, results, and conclusions, and they reviewed and verified every AI-assisted contribution. Anthropic’s Claude and Claude Code and OpenAI’s ChatGPT and Codex were used to assist with manuscript drafting and editing; with the development, debugging, and review of research software and analysis scripts; and with checking reported numbers against the recorded benchmark data, including runtime aggregation, speedup calculations, and consistency among the manuscript, figures, tables, and underlying data.

## Supplementary Information

Extended Figures and Supplementary Information are available for this paper.

## Acknowledgements

The authors gratefully acknowledge All of Us participants for their contributions, without whom this research would not have been possible. The authors thank Andrew Clark and Can Firtina for their feedback on the manuscript. In addition, we thank the National Institutes of Health All of Us Research Program for making available the participant data examined in this study. This study used data from the All of Us Research Program Controlled Tier Dataset CDRv8, available to authorized users on the Researcher Workbench. This material is based upon work supported by the National Science Foundation under Grant IIS-2435801. This research used resources from the National Energy Research Scientific Computing Center, a DOE Office of Science User Facility supported by the Office of Science of the U.S. Department of Energy under Contract No. DE-AC02-05CH11231, using NERSC award ASCR-ERCAP0030076. This work used DeltaAI at the National Center for Supercomputing Applications (NCSA) through allocation CIS251351 from the Advanced Cyberinfrastructure Coordination Ecosystem: Services & Support (ACCESS) program, which is supported by U.S. National Science Foundation grants #2138259, #2138286, #2138307, #2137603, and #2138296.

## Declarations

### Ethics Approval and Consent to Participate

Not applicable. All analyses use de-identified data accessed under approved data-use agreements with the All of Us Research Program and the publicly available 1000 Genomes Project. This study analyzed de-identified human genomic data and generated no new human-subjects data. The analyses of *All of Us* data were conducted on the *All of Us* Researcher Workbench under an approved Data Use and Registration Agreement, in accordance with the *All of Us* Research Program’s data-use policies and controlled-tier access requirements; the *All of Us* Research Program obtained ethical approval and participant informed consent for the collection and research use of these data, as described by the program [1]. The 1000 Genomes samples were collected with informed consent for open-access research use by the original studies [33].

### Consent for Publication

Not applicable.

### Data Availability

The individual-level *All of Us* data are available to authorized researchers through the *All of Us* Researcher Workbench under its controlled tier access framework and cannot be redistributed by the authors; access is granted by the *All of Us* Research Program following registration and completion of its data use requirements. The 1000 Genomes data are publicly available [33]. The `.vcf.gz` files can be downloaded from https://ftp.1000genomes.ebi.ac.uk/vol1/ftp/data_collections/1000G_2504_high_coverage/working/20220422_3202_phased_SNV_INDEL_SV/. The simulated 200,000- and 500,000-individual cohorts can be downloaded from https://portal.nersc.gov/cfs/m4341/mikado/, which includes the GRG files and the PLINK files for the last four chromosomes of the 200k cohort. The analyses were conducted within the *All of Us* Researcher Workbench in a controlled-tier workspace (ID `test-workspace-grg-gg-s-pod`, UUID 8680c880-37a5-471c-b5e5-1b965ce196cd), provisioned in resource region `us-central1` on pod `user-pod-gguidi-0ef9`.

## Materials Availability

Not applicable.

## Code Availability

The CPU/CUDA reproduction repositories are consolidated at https://github.com/CornellHPC/Mikado/releases/tag/v0.2. This includes pinned versions of `grgl`, `grapp`, and `grg-spmv`, which are the three packages necessary to run Mikado and baseline grgl on the applications. Dockerfiles and benchmark scripts are also provided. ROCm support is available in GRG-SpMV ([commit d55aab8](https://github.com/CornellHPC/grapp/tree/fb4a89f87e463c0d69882c1ff61292b4aad3abfd)) and grapp (commit `fb4a89f`). The installation and API guidance are linked at https://github.com/CornellHPC/Mikado/tree/rocm-support.

## Author Contributions

Y.L. and Q.S. designed and implemented the graph-first and SpTRSV-based GPU approaches; D.D. and X.W. contributed the GRG construction toolkit and integration; Q.S. designed the BOLT-LMM-inf benchmark and statistical validation; G.G. conceptualized the methodology, supervised the HPC and sparse-linear-algebra design; S.M. and X.W. supervised the population-genetics applications; Y.L., Q.S., X.W., and G.G. wrote the manuscript with input from the other authors.

## Supplementary Information

**Extended Data Fig. 1.**
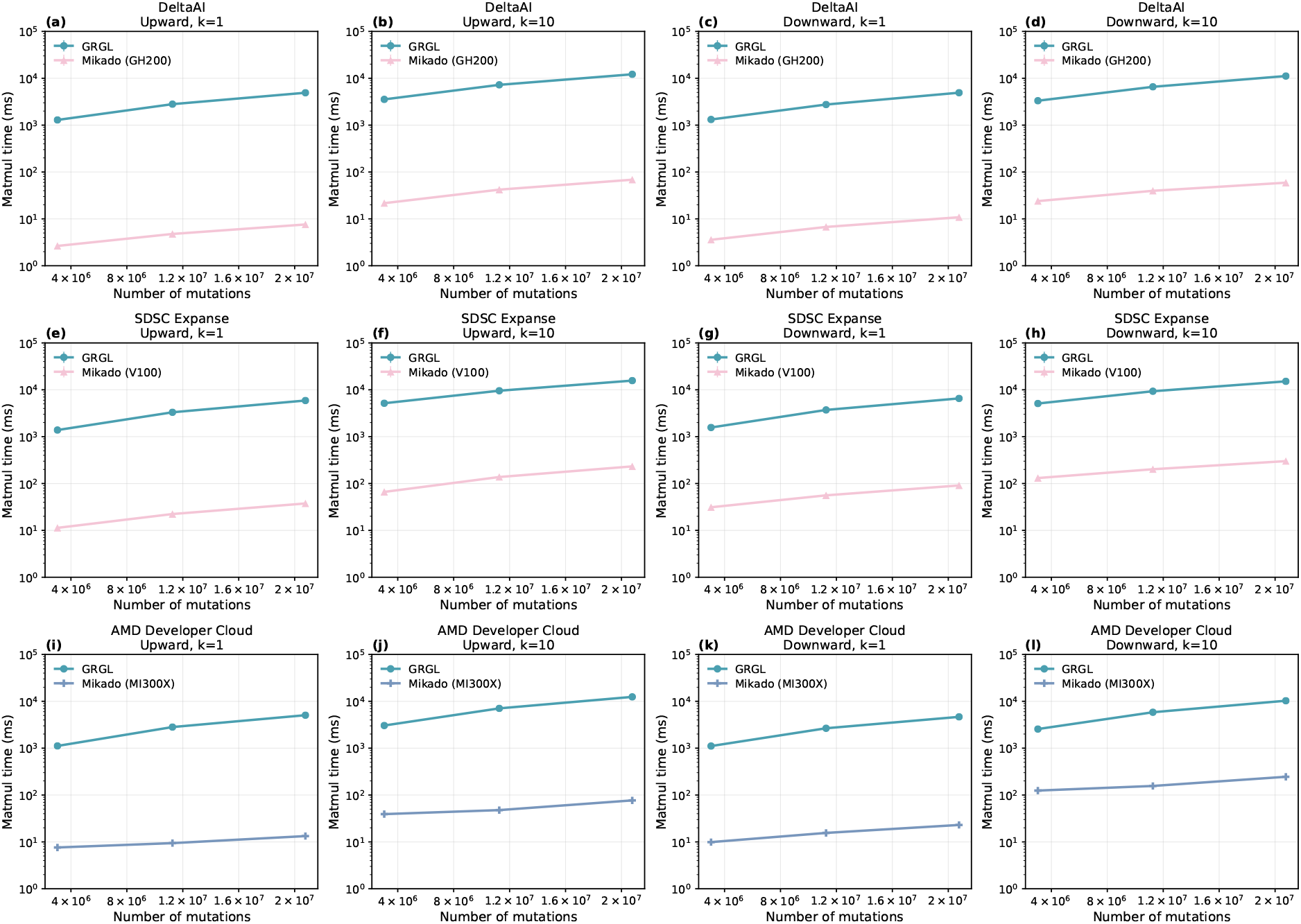
Kernel Performance on More Machines. **M****ikado** **GPU performance on DeltaAI, SDSC Expanse, and AMD Developer Cloud**. The kernel benchmark uses chromosomes 1, 11, and 21 from the simulated 500K dataset; speedups are geometric means over the three chromosomes and both directions relative to single-threaded GRGL on the same node. The CUDA code runs unchanged on the two NVIDIA generations. **(a–d)** Performance on DeltaAI, with a Grace (Arm Neoverse V2) CPU and the Hopper GPU of a GH200 superchip; GPU speedups average 483× at *k* = 1 and 167× at *k* = 10. **(e–h)** Performance on SDSC Expanse, with Intel Xeon Gold 6248 CPUs and a V100 GPU; GPU speedups average 94× at *k* = 1 and 57× at *k* = 10. **(i–l)** Performance on AMD Developer Cloud, with Intel Xeon Platinum 8568Y+ CPUs and an AMD Instinct MI300X GPU; GPU speedups average 200× at *k* = 1 and 63× at *k* = 10.

**Extended Data Fig. 2.**
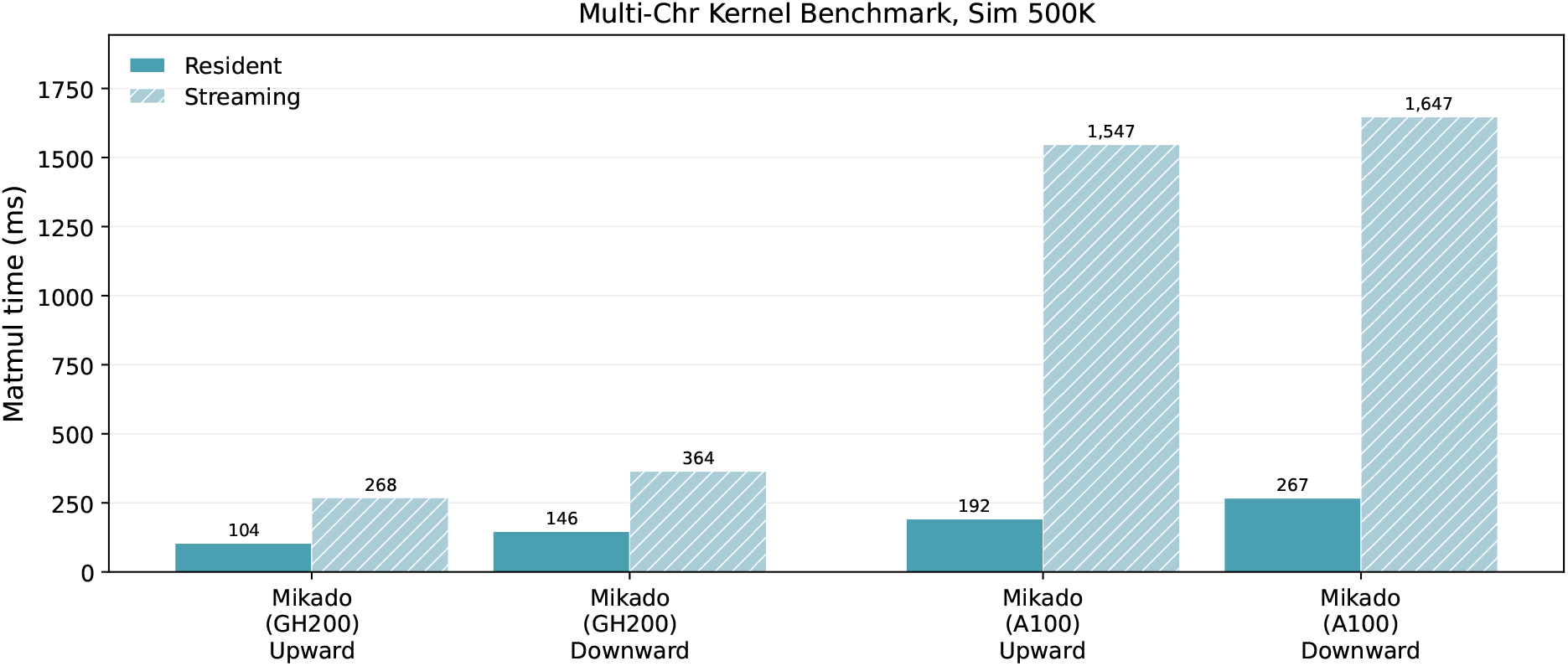
Streaming Backend Performance. **Streaming Backend Comparison**. By default, all GRGs reside in GPU memory (resident mode). For GRGs too large to fit in device memory, Mikado streams blocks on demand from host memory with double buffering, so one block is processed while the next is in transit (streaming mode). This greatly reduces GPU memory requirements, at the cost of being bound by host-to-device bandwidth rather than compute. The multi-chromosome kernel benchmarks were run on DeltaAI (one GH200 superchip) and Perlmutter (one A100 80 GB for both modes) with the simulated 500K dataset, in both resident and forced-streaming modes, to isolate the cost of streaming. On the A100, connected over PCIe 4.0 at 32 GB/s, forcing streaming incurs a slowdown exceeding 6× . On the GH200 superchip, which pairs a 72-core Grace CPU with an H100 GPU over a 900 GB/s (bidirectional) NVLink-C2C interface, the slowdown falls below 3×.

**Extended Data Fig. 3.**
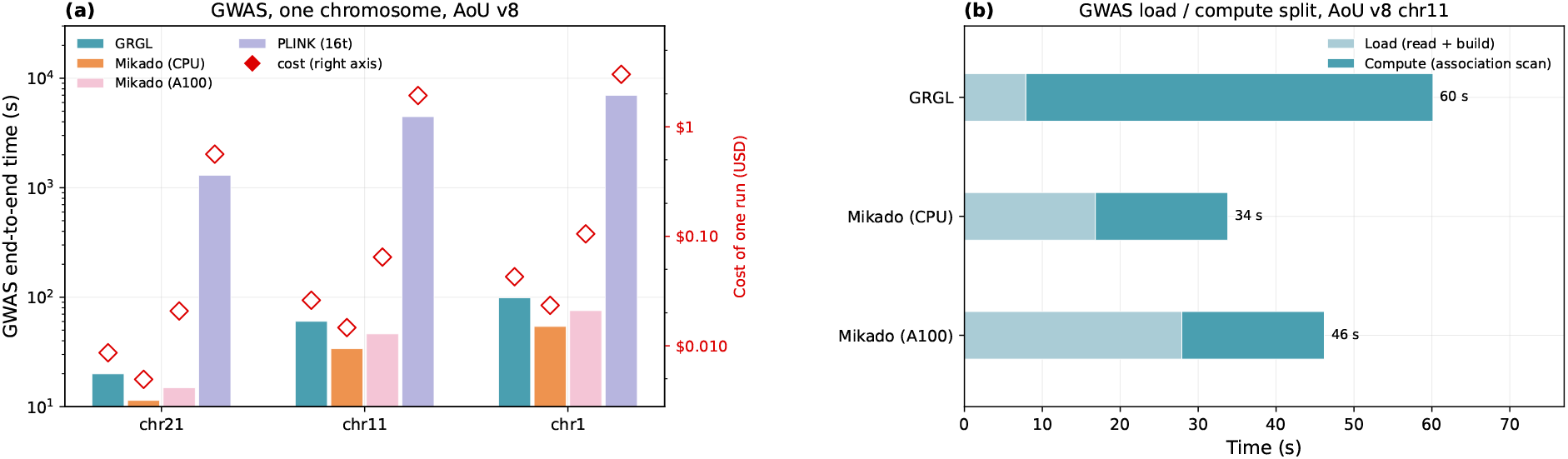
Genome-Wide Association Study (GWAS) Runtime and Cost. **Runtime and Cost of GWAS**. GWAS tests each variant for association with a phenotype, conditional on covariates; for a quantitative phenotype, the core per-variant computation reduces to a single *G*^⊤^*w* product followed by statistical post-processing. Runtime is therefore dominated by chromosome loading rather than by the matrix–vector kernel, placing GWAS at the low end of the workload intensity range described in the main text. **(a)** A single-chromosome GWAS runs on *All of Us* v8. On chromosome 11, Mikado CPU completes GWAS end to end in 34 s, compared with 4,457 s for PLINK2 and 60 s for the CPU GRG baseline, a 132× speedup over PLINK2 and 1.8× over the CPU GRG baseline; chromosomes 1 and 21 show the same pattern. Output write time is excluded for all implementations, and MKL’s `optimize` is disabled. Mikado (A100) uses one 80 GB device. **(b)** Runtime breakdown for chromosome 11, where Mikado on CPU is faster than on GPU and achieves the lowest cost.

**Extended Data Figure 4.**
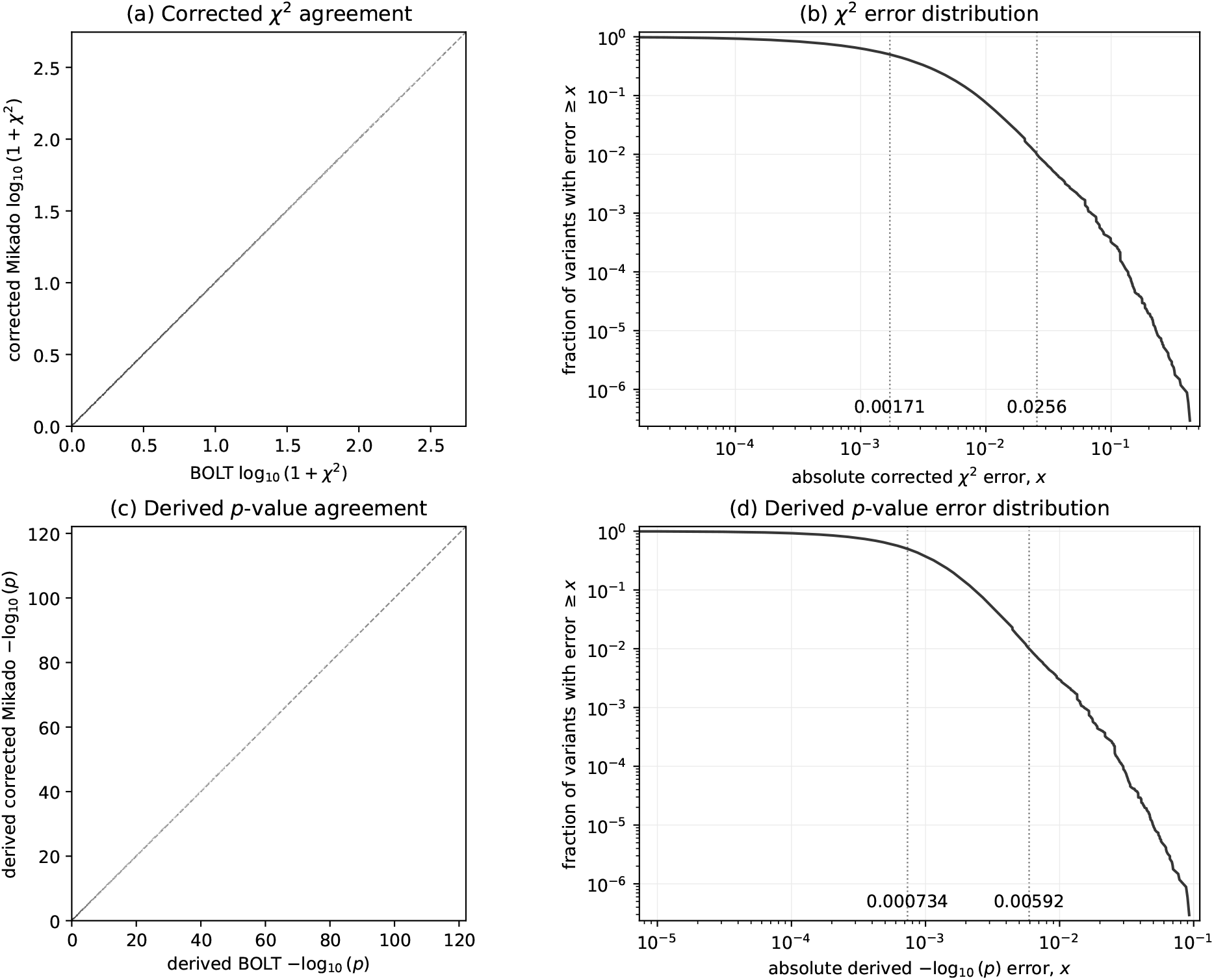
Numerical Agreement with Official BOLT-LMM-inf. **The numerical correctness of BOLT-LMM association statistics**. Comparison of official BOLT-LMM and Mikado (A100), using 40 GB, for the 200K simulation on chromosomes 19–22 after placing Mikado’s statistics on BOLT-LMM’s calibration scale. **(a)** The square-bin density of corrected *χ*^2^ statistics; the dashed line is the identity. **(b)** The empirical exceedance distribution of the absolute *χ*^2^ error. **(c**,**d)** The corresponding agreement and absolute error distributions for the derived *p* values. In both error panels **(b**,**d)**, the left and right dotted guides mark the median and 99th-percentile errors, respectively, with their values printed on the x-axis. Each output row tests one additive SNP coefficient, so its squared standardized score follows a 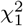 distribution. The *p*-values are recomputed from *χ*^2^ using the same 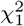 survival function because BOLT’s textual values are heavily rounded. For this diagnostic comparison only, Mikado statistics are placed on BOLT’s realized calibration scale to isolate implementation-level agreement; this rescaling is not part of the Mikado workflow and is not required in practice.

### Supplementary Note 1 | Numerical Analysis of BOLT-LMM

Our Mikado GPU implementation of BOLT-LMM differs from the official implementation in three respects: it uses a lossless GRG representation of the genotypes, algebraically equivalent kernels with a different floating-point reduction order, and NumPy PRNG rather than Boost PRNG to generate the Monte Carlo probes and calibration SNP samples. The representation is exact, and the altered reduction order affects only numerical rounding. The different PRNG changes the realized stochastic samples even when both programs receive the same seed, so two otherwise equivalent runs can produce slightly different heritability estimates and overall calibration factors.

Each of the 3,391,600 variants on chromosomes 19–22 produces outputs that match one-to-one. In the illustrated simulation with a target of *h*^2^ = 0.6, BOLT and Mikado estimate 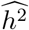 as 0.579444 and 0.577319, respectively. Importantly, Mikado does not require a BOLT-LMM reference result or any additional rescaling in practice. The heritability-recovery experiment shows that its Monte Carlo estimation and calibration procedure accurately recovers the simulated heritability targets, establishing that the method independently produces useful inference.

Extended Data Fig. 4 addresses a separate implementation question: once the known, run-specific calibration-scale difference is factored out for this diagnostic comparison, do the accelerated representation and kernels reproduce BOLT’s variant-level calculations? The independently sampled calibration factors are 1.09367 (BOLT) and 1.053465 (Mikado GPU). For this audit only, applying their ratio of 1.038164 places the statistics on a common realized calibration scale. The corrected statistics then agree essentially point for point, with a through-origin slope of 0.999955 and *R*^2^ = 0.999999871. The absolute *χ*^2^ errors have median, 99th percentile, and maximum values of 0.00171, 0.0256, and 0.425, respectively, and a relative *L*_2_ error of 0.0356%. Derived p-values yield 45,818 shared genome-wide significant calls at 5 × 10^−8^ and only 17 boundary-crossing differences among 3.39 million tests.

The recovery experiment establishes that Mikado BOLT-LMM is valid without reference correction, while the pointwise comparison shows that the accelerated computation itself introduces a negligible additional numerical error. Together, these results show that the reported speedups do not compromise inferential accuracy.

### Supplementary Note 2 | Per-Chromosome Statistics

In this section, Supplementary Tables 1–8 provide per-chromosome statistics for each dataset. The height column indicates the number of height levels for the constructed GRGs using Mikado‘s formulation.

**Supplementary Table 1.** **AoU v8 phased dataset** statistics. Cohort size: 414,830. PLINK2 is the total size of the `.pgen`/`.psam`/`.pvar` file set. PLINK2 is built only for a subset of chromosomes.

| Chr | #Mutations | VCF.gz | PLINK2 | GRG | GRG-SpMV | #Nodes | #Edges | Height |
|---|---|---|---|---|---|---|---|---|
| 1 | 69,165,511 | 238 GB | 86 GB | 8.3 GB | 16 GB | 178,677,711 | 2,026,690,586 | 30 |
| 2 | 76,695,484 | 259 GB |  | 8.7 GB | 16 GB | 182,468,494 | 2,077,649,352 | 29 |
| 3 | 63,328,899 | 218 GB |  | 7.1 GB | 13 GB | 150,127,703 | 1,692,271,935 | 29 |
| 4 | 61,362,449 | 216 GB |  | 6.9 GB | 13 GB | 145,737,061 | 1,646,949,664 | 56 |
| 5 | 57,032,792 | 196 GB |  | 6.4 GB | 12 GB | 136,152,008 | 1,536,836,594 | 30 |
| 6 | 53,288,431 | 191 GB |  | 6.1 GB | 12 GB | 128,666,201 | 1,456,900,356 | 33 |
| 7 | 50,416,022 | 175 GB |  | 6.1 GB | 12 GB | 128,581,218 | 1,488,117,888 | 44 |
| 8 | 49,585,609 | 167 GB |  | 5.6 GB | 11 GB | 117,857,735 | 1,365,867,818 | 41 |
| 9 | 39,350,131 | 133 GB |  | 5.0 GB | 9.2 GB | 107,003,860 | 1,278,132,610 | 34 |
| 10 | 42,213,334 | 149 GB |  | 5.2 GB | 9.6 GB | 111,955,468 | 1,303,030,322 | 29 |
| 11 | 43,024,023 | 149 GB | 55 GB | 5.0 GB | 9.2 GB | 106,533,797 | 1,217,490,442 | 28 |
| 12 | 41,172,758 | 144 GB |  | 5.0 GB | 9.1 GB | 106,381,765 | 1,211,686,472 | 27 |
| 13 | 30,469,204 | 108 GB |  | 3.7 GB | 6.8 GB | 79,740,553 | 903,900,867 | 28 |
| 14 | 27,959,778 | 98 GB |  | 3.5 GB | 6.4 GB | 74,262,590 | 866,463,284 | 27 |
| 15 | 25,724,763 | 89 GB |  | 3.5 GB | 6.5 GB | 76,098,461 | 916,442,328 | 28 |
| 16 | 29,179,546 | 95 GB |  | 3.8 GB | 7.0 GB | 82,276,293 | 974,608,659 | 27 |
| 17 | 24,574,646 | 84 GB |  | 3.4 GB | 6.3 GB | 75,226,753 | 886,785,072 | 27 |
| 18 | 23,870,313 | 84 GB |  | 3.1 GB | 5.6 GB | 67,546,265 | 772,505,926 | 28 |
| 19 | 18,727,782 | 66 GB |  | 2.8 GB | 5.1 GB | 61,060,691 | 728,921,352 | 28 |
| 20 | 19,247,900 | 67 GB |  | 2.6 GB | 4.8 GB | 57,416,369 | 669,478,407 | 27 |
| 21 | 11,974,519 | 41 GB | 16 GB | 1.8 GB | 3.3 GB | 37,155,828 | 487,056,362 | 26 |
| 22 | 11,973,519 | 42 GB |  | 1.9 GB | 3.5 GB | 40,213,718 | 520,556,991 | 26 |
| **Total** | 870,337,413 | 2.94 TB |  | 106 GB | 197 GB | 2,251,140,542 | 26,028,343,287 |  |

**Supplementary Table 2.** **AoU v8 phased, filtered dataset** statistics (non-SNP and multiallelic sites removed). Cohort size: 414, 830. Only chr21 (filtered) is used in the main text (Fig. 3a); chr1 and chr11 are listed for reference.

| Chr | #Mutations | PLINK2 | GRG | GRG-SpMV | #Nodes | #Edges |
|---|---|---|---|---|---|---|
| 1 | 36,917,066 | 41 GB | 6.7 GB | 12 GB | 161,731,460 | 1,717,867,968 |
| 11 | 22,395,732 | 25 GB | 4.0 GB | 7.2 GB | 95,759,938 | 1,036,625,578 |
| 21 | 5,851,262 | 7.0 GB | 1.4 GB | 2.5 GB | 32,446,275 | 387,518,340 |

**Supplementary Table 3.** **AoU v8 phased filtered dataset II** used for the BOLT-LMM experiment. Cohort size: 220,760. The sites with MAF ≥ 5 × 10^−5^ and Hardy–Weinberg equilibrium *p* ≥ 10^−12^ were retained; non-SNP and multi-allelic sites were removed; individuals were restricted to the EUR group in the ancestry file, excluding those with quality-control flags.

| Chr | #Mutations | GRG | GRG-SpMV | #Nodes | #Edges |
|---|---|---|---|---|---|
| 1 | 3,619,944 | 2.0 GB | 3.7 GB | 39,735,582 | 708,528,044 |
| 2 | 3,913,024 | 2.0 GB | 3.8 GB | 40,835,120 | 732,770,801 |
| 3 | 3,226,487 | 1.7 GB | 3.1 GB | 34,118,223 | 601,630,061 |
| 4 | 3,121,749 | 1.7 GB | 3.1 GB | 33,165,115 | 592,890,256 |
| 5 | 2,937,757 | 1.5 GB | 2.9 GB | 31,012,135 | 551,858,873 |
| 6 | 2,792,088 | 1.5 GB | 2.8 GB | 29,473,889 | 526,576,298 |
| 7 | 2,593,774 | 1.5 GB | 2.7 GB | 28,840,247 | 524,975,255 |
| 8 | 2,428,240 | 1.3 GB | 2.5 GB | 25,456,569 | 474,583,211 |
| 9 | 1,943,053 | 1.2 GB | 2.2 GB | 22,420,483 | 431,122,970 |
| 10 | 2,206,388 | 1.3 GB | 2.4 GB | 24,887,211 | 456,271,903 |
| 11 | 2,194,556 | 1.2 GB | 2.3 GB | 24,041,465 | 433,277,839 |
| 12 | 2,141,418 | 1.2 GB | 2.2 GB | 23,866,859 | 431,163,245 |
| 13 | 1,586,417 | 883 MB | 1.7 GB | 17,800,704 | 321,096,982 |
| 14 | 1,456,131 | 803 MB | 1.6 GB | 15,924,201 | 297,762,981 |
| 15 | 1,313,544 | 776 MB | 1.5 GB | 15,174,935 | 294,099,974 |
| 16 | 1,362,787 | 830 MB | 1.6 GB | 16,333,812 | 313,652,422 |
| 17 | 1,311,961 | 793 MB | 1.5 GB | 15,707,133 | 296,683,801 |
| 18 | 1,254,076 | 737 MB | 1.4 GB | 14,802,780 | 270,155,968 |
| 19 | 973,237 | 631 MB | 1.2 GB | 12,382,695 | 239,630,388 |
| 20 | 1,026,488 | 620 MB | 1.2 GB | 12,340,651 | 229,143,277 |
| 21 | 603,268 | 385 MB | 752 MB | 7,175,678 | 151,465,316 |
| 22 | 620,556 | 413 MB | 811 MB | 7,583,591 | 164,855,247 |
| **Total** | 44,626,943 | 25 GB | 47 GB | 493,079,078 | 9,044,195,112 |

**Supplementary Table 4.**
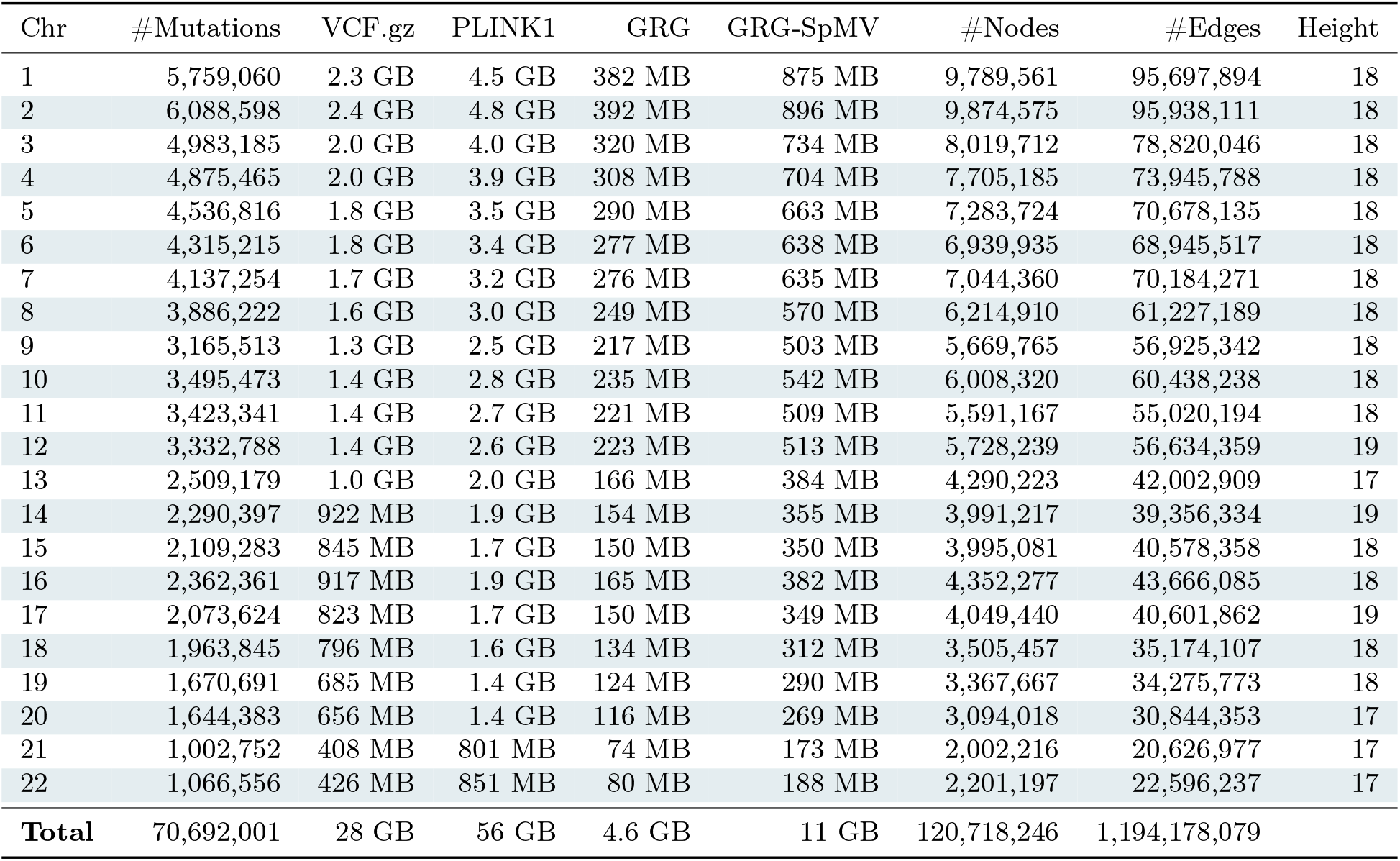
**1000 Genomes dataset** statistics. Cohort size: 3, 202. PLINK1 is the total size of the `.bed`/`.bim`/`.fam` fileset.

| Chr | #Mutations | VCF.gz | PLINK1 | GRG | GRG-SpMV | #Nodes | #Edges | Height |
|---|---|---|---|---|---|---|---|---|
| 1 | 5,759,060 | 2.3 GB | 4.5 GB | 382 MB | 875 MB | 9,789,561 | 95,697,894 | 18 |
| 2 | 6,088,598 | 2.4 GB | 4.8 GB | 392 MB | 896 MB | 9,874,575 | 95,938,111 | 18 |
| 3 | 4,983,185 | 2.0 GB | 4.0 GB | 320 MB | 734 MB | 8,019,712 | 78,820,046 | 18 |
| 4 | 4,875,465 | 2.0 GB | 3.9 GB | 308 MB | 704 MB | 7,705,185 | 73,945,788 | 18 |
| 5 | 4,536,816 | 1.8 GB | 3.5 GB | 290 MB | 663 MB | 7,283,724 | 70,678,135 | 18 |
| 6 | 4,315,215 | 1.8 GB | 3.4 GB | 277 MB | 638 MB | 6,939,935 | 68,945,517 | 18 |
| 7 | 4,137,254 | 1.7 GB | 3.2 GB | 276 MB | 635 MB | 7,044,360 | 70,184,271 | 18 |
| 8 | 3,886,222 | 1.6 GB | 3.0 GB | 249 MB | 570 MB | 6,214,910 | 61,227,189 | 18 |
| 9 | 3,165,513 | 1.3 GB | 2.5 GB | 217 MB | 503 MB | 5,669,765 | 56,925,342 | 18 |
| 10 | 3,495,473 | 1.4 GB | 2.8 GB | 235 MB | 542 MB | 6,008,320 | 60,438,238 | 18 |
| 11 | 3,423,341 | 1.4 GB | 2.7 GB | 221 MB | 509 MB | 5,591,167 | 55,020,194 | 18 |
| 12 | 3,332,788 | 1.4 GB | 2.6 GB | 223 MB | 513 MB | 5,728,239 | 56,634,359 | 19 |
| 13 | 2,509,179 | 1.0 GB | 2.0 GB | 166 MB | 384 MB | 4,290,223 | 42,002,909 | 17 |
| 14 | 2,290,397 | 922 MB | 1.9 GB | 154 MB | 355 MB | 3,991,217 | 39,356,334 | 19 |
| 15 | 2,109,283 | 845 MB | 1.7 GB | 150 MB | 350 MB | 3,995,081 | 40,578,358 | 18 |
| 16 | 2,362,361 | 917 MB | 1.9 GB | 165 MB | 382 MB | 4,352,277 | 43,666,085 | 18 |
| 17 | 2,073,624 | 823 MB | 1.7 GB | 150 MB | 349 MB | 4,049,440 | 40,601,862 | 19 |
| 18 | 1,963,845 | 796 MB | 1.6 GB | 134 MB | 312 MB | 3,505,457 | 35,174,107 | 18 |
| 19 | 1,670,691 | 685 MB | 1.4 GB | 124 MB | 290 MB | 3,367,667 | 34,275,773 | 18 |
| 20 | 1,644,383 | 656 MB | 1.4 GB | 116 MB | 269 MB | 3,094,018 | 30,844,353 | 17 |
| 21 | 1,002,752 | 408 MB | 801 MB | 74 MB | 173 MB | 2,002,216 | 20,626,977 | 17 |
| 22 | 1,066,556 | 426 MB | 851 MB | 80 MB | 188 MB | 2,201,197 | 22,596,237 | 17 |
| **Total** | 70,692,001 | 28 GB | 56 GB | 4.6 GB | 11 GB | 120,718,246 | 1,194,178,079 |  |

**Supplementary Table 5.**
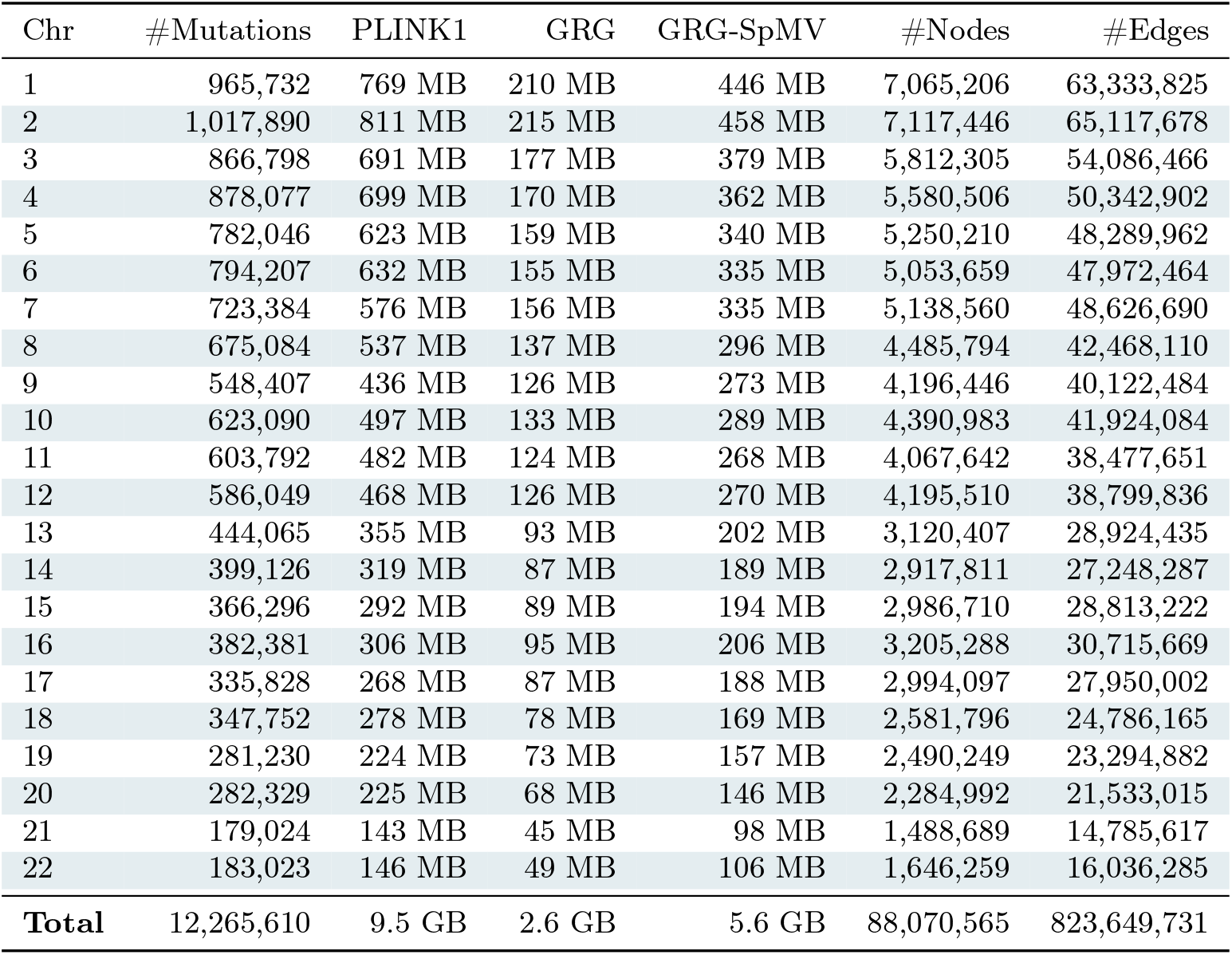
**1000 Genomes filtered dataset** statistics. Cohort size: 3, 202. The sites with MAF ≥ 10^−2^ are removed, multiallelic sites, and non-SNP sites are removed.

| Chr | #Mutations | PLINK1 | GRG | GRG-SpMV | #Nodes | #Edges |
|---|---|---|---|---|---|---|
| 1 | 965,732 | 769 MB | 210 MB | 446 MB | 7,065,206 | 63,333,825 |
| 2 | 1,017,890 | 811 MB | 215 MB | 458 MB | 7,117,446 | 65,117,678 |
| 3 | 866,798 | 691 MB | 177 MB | 379 MB | 5,812,305 | 54,086,466 |
| 4 | 878,077 | 699 MB | 170 MB | 362 MB | 5,580,506 | 50,342,902 |
| 5 | 782,046 | 623 MB | 159 MB | 340 MB | 5,250,210 | 48,289,962 |
| 6 | 794,207 | 632 MB | 155 MB | 335 MB | 5,053,659 | 47,972,464 |
| 7 | 723,384 | 576 MB | 156 MB | 335 MB | 5,138,560 | 48,626,690 |
| 8 | 675,084 | 537 MB | 137 MB | 296 MB | 4,485,794 | 42,468,110 |
| 9 | 548,407 | 436 MB | 126 MB | 273 MB | 4,196,446 | 40,122,484 |
| 10 | 623,090 | 497 MB | 133 MB | 289 MB | 4,390,983 | 41,924,084 |
| 11 | 603,792 | 482 MB | 124 MB | 268 MB | 4,067,642 | 38,477,651 |
| 12 | 586,049 | 468 MB | 126 MB | 270 MB | 4,195,510 | 38,799,836 |
| 13 | 444,065 | 355 MB | 93 MB | 202 MB | 3,120,407 | 28,924,435 |
| 14 | 399,126 | 319 MB | 87 MB | 189 MB | 2,917,811 | 27,248,287 |
| 15 | 366,296 | 292 MB | 89 MB | 194 MB | 2,986,710 | 28,813,222 |
| 16 | 382,381 | 306 MB | 95 MB | 206 MB | 3,205,288 | 30,715,669 |
| 17 | 335,828 | 268 MB | 87 MB | 188 MB | 2,994,097 | 27,950,002 |
| 18 | 347,752 | 278 MB | 78 MB | 169 MB | 2,581,796 | 24,786,165 |
| 19 | 281,230 | 224 MB | 73 MB | 157 MB | 2,490,249 | 23,294,882 |
| 20 | 282,329 | 225 MB | 68 MB | 146 MB | 2,284,992 | 21,533,015 |
| 21 | 179,024 | 143 MB | 45 MB | 98 MB | 1,488,689 | 14,785,617 |
| 22 | 183,023 | 146 MB | 49 MB | 106 MB | 1,646,259 | 16,036,285 |
| **Total** | 12,265,610 | 9.5 GB | 2.6 GB | 5.6 GB | 88,070,565 | 823,649,731 |

**Supplementary Table 6.** **Simulated 200k dataset** statistics. Cohort size: 200, 000.

| Chr | #Mutations | GRG | GRG-SpMV | #Nodes | #Edges | Height |
|---|---|---|---|---|---|---|
| 1 | 17,083,232 | 1.5 GB | 2.9 GB | 32,951,213 | 309,717,578 | 63 |
| 2 | 16,637,528 | 1.4 GB | 2.8 GB | 31,484,527 | 284,487,370 | 305 |
| 3 | 13,625,360 | 1.2 GB | 2.3 GB | 26,039,836 | 243,055,123 | 64 |
| 4 | 13,058,115 | 1.1 GB | 2.2 GB | 25,327,665 | 236,364,913 | 61 |
| 5 | 12,474,779 | 1.1 GB | 2.1 GB | 23,971,215 | 225,747,613 | 58 |
| 6 | 11,730,038 | 1011 MB | 2.0 GB | 22,587,449 | 213,923,794 | 68 |
| 7 | 10,960,584 | 970 MB | 1.9 GB | 21,690,881 | 210,715,273 | 100 |
| 8 | 9,953,740 | 882 MB | 1.8 GB | 19,561,095 | 193,397,364 | 89 |
| 9 | 9,500,000 | 847 MB | 1.7 GB | 18,533,938 | 191,235,205 | 97 |
| 10 | 9,196,528 | 869 MB | 1.7 GB | 19,537,600 | 198,856,189 | 70 |
| 11 | 9,265,101 | 820 MB | 1.6 GB | 18,130,386 | 180,033,855 | 78 |
| 12 | 9,158,771 | 856 MB | 1.7 GB | 19,173,025 | 195,939,104 | 56 |
| 13 | 6,598,679 | 630 MB | 1.3 GB | 13,834,971 | 148,396,050 | 64 |
| 14 | 6,034,574 | 596 MB | 1.2 GB | 12,972,483 | 145,908,504 | 67 |
| 15 | 5,656,206 | 629 MB | 1.3 GB | 13,865,922 | 167,617,037 | 52 |
| 16 | 6,224,437 | 629 MB | 1.3 GB | 13,628,263 | 159,058,378 | 99 |
| 17 | 5,714,000 | 601 MB | 1.2 GB | 13,119,720 | 154,601,782 | 54 |
| 18 | 5,523,349 | 559 MB | 1.1 GB | 12,139,949 | 140,091,898 | 75 |
| 19 | 4,010,942 | 479 MB | 964 MB | 10,311,874 | 135,031,919 | 64 |
| 20 | 4,415,250 | 486 MB | 969 MB | 10,549,213 | 129,168,016 | 52 |
| 21 | 2,502,559 | 306 MB | 616 MB | 6,088,637 | 91,467,812 | 50 |
| 22 | 2,441,331 | 332 MB | 676 MB | 6,696,671 | 104,365,793 | 46 |
| **Total** | 191,765,103 | 18 GB | 35 GB | 392,196,533 | 4,059,180,570 |  |

**Supplementary Table 7.** **Simulated 200k filtered dataset** statistics. Cohort size: 200, 000. The sites with MAF *<* 10^−4^ and multiallelic sites are removed. PLINK1 is the total size of the `.bed`/`.bim`/`.fam` fileset.

| Chr | #Mutations | PLINK1 | GRG | GRG-SpMV | #Nodes | #Edges |
|---|---|---|---|---|---|---|
| 19 | 1,016,718 | 48 GB | 377 MB | 771 MB | 9,153,249 | 130,648,570 |
| 20 | 1,117,819 | 53 GB | 376 MB | 759 MB | 9,314,525 | 124,934,949 |
| 21 | 638,234 | 30 GB | 242 MB | 496 MB | 5,289,198 | 89,054,606 |
| 22 | 618,829 | 29 GB | 268 MB | 556 MB | 5,912,762 | 101,491,930 |

**Supplementary Table 8.** **Simulated 500k dataset** statistics. Cohort size: 500, 000.

| Chr | #Mutations | GRG | GRG-SpMV | #Nodes | #Edges | Height |
|---|---|---|---|---|---|---|
| 1 | 20,769,289 | 2.1 GB | 4.0 GB | 46,026,894 | 468,916,211 | 66 |
| 2 | 20,205,851 | 2.0 GB | 3.8 GB | 43,658,912 | 427,330,611 | 62 |
| 3 | 16,541,902 | 1.7 GB | 3.2 GB | 36,318,030 | 375,881,430 | 69 |
| 4 | 15,865,658 | 1.6 GB | 3.1 GB | 35,148,353 | 368,334,094 | 54 |
| 5 | 15,148,360 | 1.5 GB | 2.9 GB | 33,253,490 | 350,317,217 | 65 |
| 6 | 14,248,401 | 1.5 GB | 2.8 GB | 31,425,705 | 334,271,994 | 67 |
| 7 | 13,296,591 | 1.4 GB | 2.7 GB | 30,071,822 | 329,546,973 | 97 |
| 8 | 12,106,619 | 1.3 GB | 2.5 GB | 27,077,962 | 304,925,641 | 51 |
| 9 | 11,593,486 | 1.3 GB | 2.4 GB | 26,007,829 | 306,957,558 | 87 |
| 10 | 11,173,189 | 1.3 GB | 2.4 GB | 27,089,894 | 313,091,390 | 88 |
| 11 | 11,259,696 | 1.2 GB | 2.3 GB | 25,315,430 | 290,486,081 | 68 |
| 12 | 11,127,551 | 1.3 GB | 2.4 GB | 26,633,545 | 309,080,942 | 46 |
| 13 | 8,009,106 | 937 MB | 1.8 GB | 19,177,193 | 246,468,052 | 52 |
| 14 | 7,325,817 | 900 MB | 1.8 GB | 18,092,477 | 248,323,688 | 49 |
| 15 | 6,847,885 | 952 MB | 1.9 GB | 19,241,764 | 281,814,674 | 55 |
| 16 | 7,541,581 | 941 MB | 1.9 GB | 19,038,920 | 262,937,910 | 82 |
| 17 | 6,947,200 | 914 MB | 1.8 GB | 18,340,394 | 263,842,135 | 61 |
| 18 | 6,709,681 | 857 MB | 1.7 GB | 17,074,882 | 242,590,328 | 67 |
| 19 | 4,870,210 | 745 MB | 1.5 GB | 14,367,819 | 234,885,598 | 53 |
| 20 | 5,371,399 | 760 MB | 1.5 GB | 14,815,210 | 228,835,097 | 51 |
| 21 | 3,034,975 | 519 MB | 1.1 GB | 8,673,158 | 180,180,643 | 61 |
| 22 | 2,967,369 | 557 MB | 1.2 GB | 9,483,299 | 199,325,274 | 45 |
| **Total** | 232,961,816 | 26 GB | 51 GB | 546,332,982 | 6,568,343,541 |  |

### Supplementary Note 3 | Mikado Artifacts

The CPU and CUDA artifacts are available at https://github.com/CornellHPC/Mikado/releases/tag/v0.2. This repository contains:

1. README file.
2. Pinned version of GRGL (Genotype Representation Graph Library), which provides core functionalities such as constructing GRGs, as well as the baseline CPU backend.
3. Pinned version of `grapp`, which provides application implementations based on GRG.
4. Pinned version of GRG-SpMV, which provides the core functionalities of the Mikado backend for `grapp`.
5. `docker/` directory containing the dockerfiles for building the images.
6. `benchmark/` directory containing the benchmark scripts and config files.
7. `code/` directory containing auxiliary code files for evaluation purposes.

Docker-based setup is recommended, though manual installation is also possible. The following examples set up and run Mikado with `podman-hpc` on the Perlmutter system; commands using `docker` are similar.

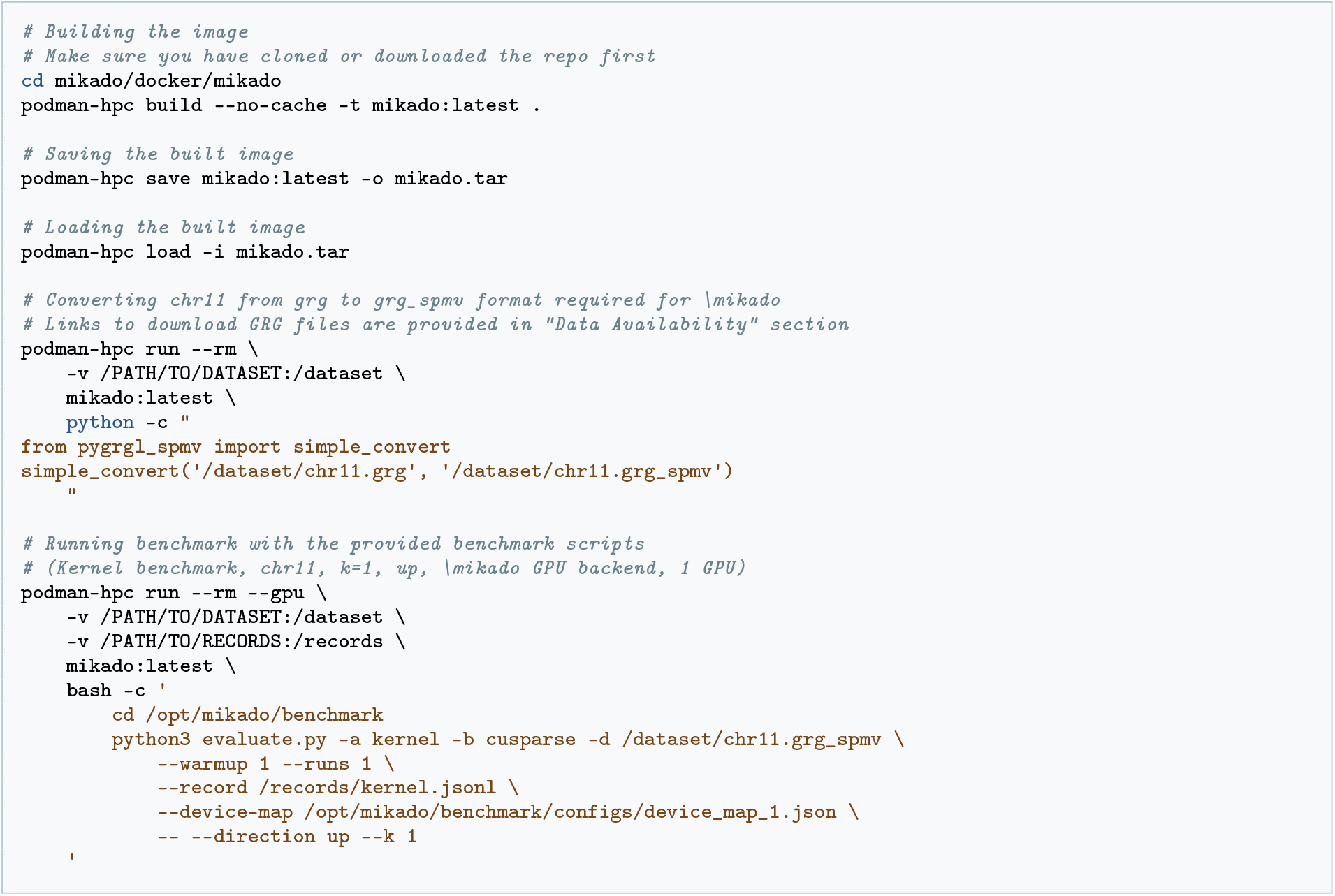

The README contains detailed instructions on how to install manually, build GRG, convert GRG, and run experiments with the benchmark scripts. The simulated datasets in `.grg` format can be downloaded from https://portal.nersc.gov/cfs/m4341/mikado/. To build GRGs from `vcf.gz` datasets, refer to the GRGL documentation at https://grgl.readthedocs.io/en/stable/construct.html.

#### ROCm Backend and Portability to AMD GPUs

ROCm support is available in GRG-SpMV (commit `d55aab8`) and `grapp` (commit `fb4a89f`), with installation and API guidance at https://github.com/CornellHPC/Mikado/tree/rocm-support. The port preserves the sparse formulation and shared application interfaces, adapting GPU runtime operations and sparse primitives to HIP and rocSPARSE. It supports resident execution on a single GPU, including multiple GRGs, but does not support multi-GPU execution or host-memory streaming. The ROCm backend was evaluated on a single AMD Instinct MI300X on AMD Developer Cloud (Supplementary Table 15) using the pip ROCm SDK 10.1.0, rocSPARSE 5.1.0, and HIP-enabled CuPy; the complete software environment, execution parameters, and input hashes are recorded with the benchmark data. Every AMD experiment used simulated data with float64 arithmetic. Extended Data Fig. 1 compares kernel performance with single-threaded GRGL on the same host, and Supplementary Table 9 reports ten-component PCA on the unfiltered 200K and 500K cohorts and BOLT-LMM-inf on filtered chromosomes 19–22 of the 200K cohort, without covariates. Repetitions, aggregation, and numerical verification follow Statistics and Reproducibility in Methods; each of the six measured AMD PCA runs met the shared residual criterion. The BOLT phenotype was regenerated using the CUDA simulation recipe without verifying its identity against the earlier CUDA phenotype, so the AMD BOLT experiment provides only runtime evidence, without a new paired accuracy comparison with official BOLT-LMM. Cross-platform runtime differences also reflect host hardware and GPU count.

**Supplementary Table 9.**
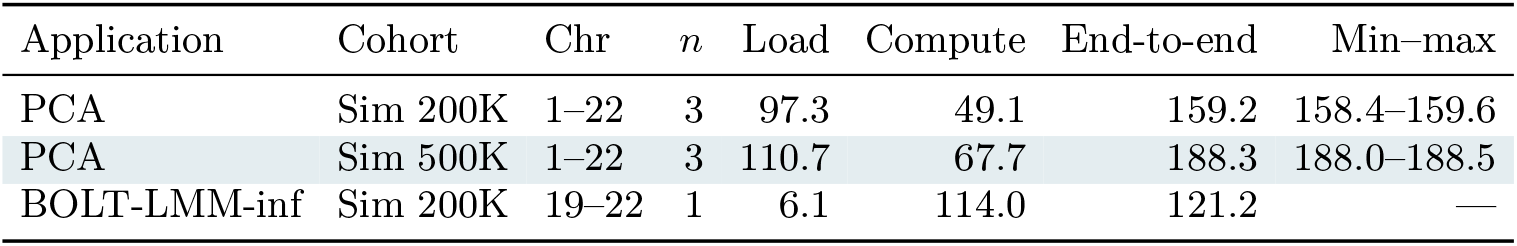
**M****ikado** **(AMD GPU, MI300X)** application runtimes in seconds. Times are process means; *n* counts measured processes, with one warm-up process per case excluded. The min–max range spans the measured end-to-end times; BOLT has a single measurement and no spread estimate. The end-to-end time includes setup beyond the load and compute timers; BOLT-LMM output writing is not included.

| Application | Cohort | Chr | *n* | Load | Compute | End-to-end | Min–max |
|---|---|---|---|---|---|---|---|
| PCA | Sim 200K | 1–22 | 3 | 97.3 | 49.1 | 159.2 | 158.4–159.6 |
| PCA | Sim 500K | 1–22 | 3 | 110.7 | 67.7 | 188.3 | 188.0–188.5 |
| BOLT-LMM-inf | Sim 200K | 19–22 | 1 | 6.1 | 114.0 | 121.2 | — |

### Supplementary Note 4 | Parameters

Each backend in the benchmark suite may require specific parameters that affect performance and, in rare cases, accuracy. This section describes these parameter settings in detail. The main evaluation script (`evaluate.py`) is called with these parameters.

#### Mikado CPU (MKL)

1. `--mkl-threads <N>`: number of MKL threads per chromosome.
2. `--optimize`: enable MKL optimization flags.

The settings for these options are shown in Supplementary Table 10.

**Supplementary Table 10.** **M****ikado** **CPU (MKL)** parameters.

| Application | Reference Figure | --mkl-threads | --optimize |
|---|---|---|---|
| Kernel | Fig. 2 (c–f) | 1 | Yes |
| Kernel | Fig. 2 (g–h) | Varies | Varies |
| PCA | Fig. 3 (a,c) | 16 | Yes |
| PCA | Fig. 3 (b) | 1 | No |
| PCA | Fig. 3 (d) | 1 | Varies |
| BOLT-LMM | Fig. 4 | 1 | No |
| GWAS | Extended Fig. 3 | 16 | No |

#### Mikado GPU cuSPARSE/rocSPARSE

1. `--native`: use CuPy arrays, GPU-to-GPU communication, and CuPy-based PCA; set for all experiments except GWAS (Extended Fig. 3).
2. `--capture`: enable CUDA or HIP graph capture; set for all GPU experiments.
3. `--device-map <FILE>`: control the CUDA mapping of chromosomes onto devices, used to set the number of GPUs in benchmarks. `device_map_1.json` is used for all single-chromosome CUDA experiments.
4. `--force-spmm`: force *k* = 2 when computing *k* = 1 to avoid a cuSPARSE precision bug for SpMV in earlier CUDA versions; **not** set for any experiment.
5. `--tol-record <FILE>`: record file used to align stopping criteria for CuPy-based PCA runs; set to the corresponding record file from GRGL runs for CUDA PCA experiments.
6. `--no-allow-residency --vram-budget-mb <N>`: CUDA-only options used together to set a GPU memory budget and force streaming mode; set only for the Streaming Backend Comparison (Extended Fig. 2), for which the GPU memory budget is 8000 MB.

The settings for the device-map option in multi-chromosome experiments are shown in Supplementary Table 11. ROCm implementation details and application experiments are described in Supplementary Note 3; solver settings for PCA and BOLT-LMM-inf, including stopping tolerances, are given under Statistics and Reproducibility in Methods.

**Supplementary Table 11.** **M****ikado** **GPU (cuSPARSE)** parameters.

| Application | Figure | --device-map (device_map_{n}.json) |
|---|---|---|
| PCA | Fig. 3 (b) | 4 |
| BOLT-LMM | Fig. 4 | 1 |

#### PLINK2

`--threads <N>`: the number of threads.

The settings for the thread option are shown in Supplementary Table 12. Below are the commands the benchmark scripts use to invoke PLINK2.

*Invoke command for PLINK2 PCA*:

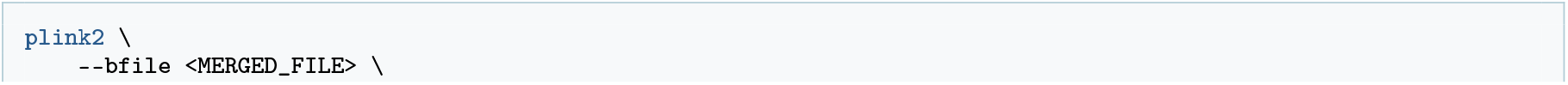

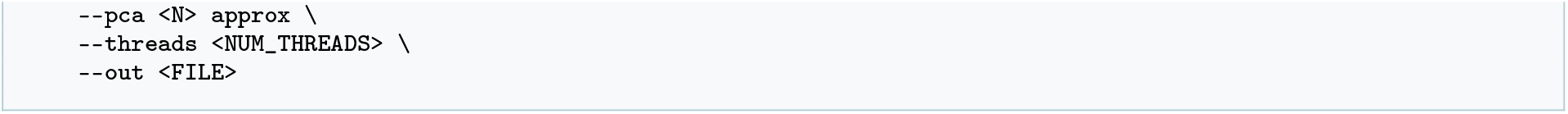

*Invoke command for PLINK2 GWAS*:

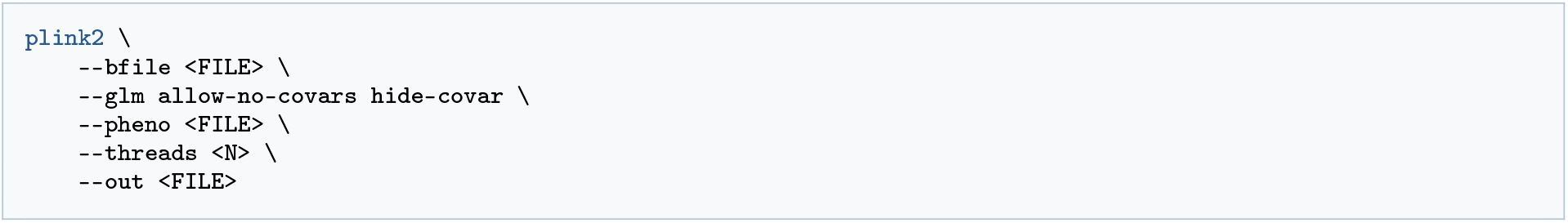

**Supplementary Table 12.** **PLINK2** parameters.

| Application | Figure | --threads |
|---|---|---|
| PCA | Fig. 3 (a) | 16 |
| PCA | Fig. 3 (b) | 64 |
| GWAS | Extended Fig. 3 (a) | 16 |

#### BOLT-LMM

1. `--threads <N>`: the number of threads.

The settings for the thread option are shown in Supplementary Table 13. Below are the commands the benchmark scripts use to invoke BOLT-LMM.

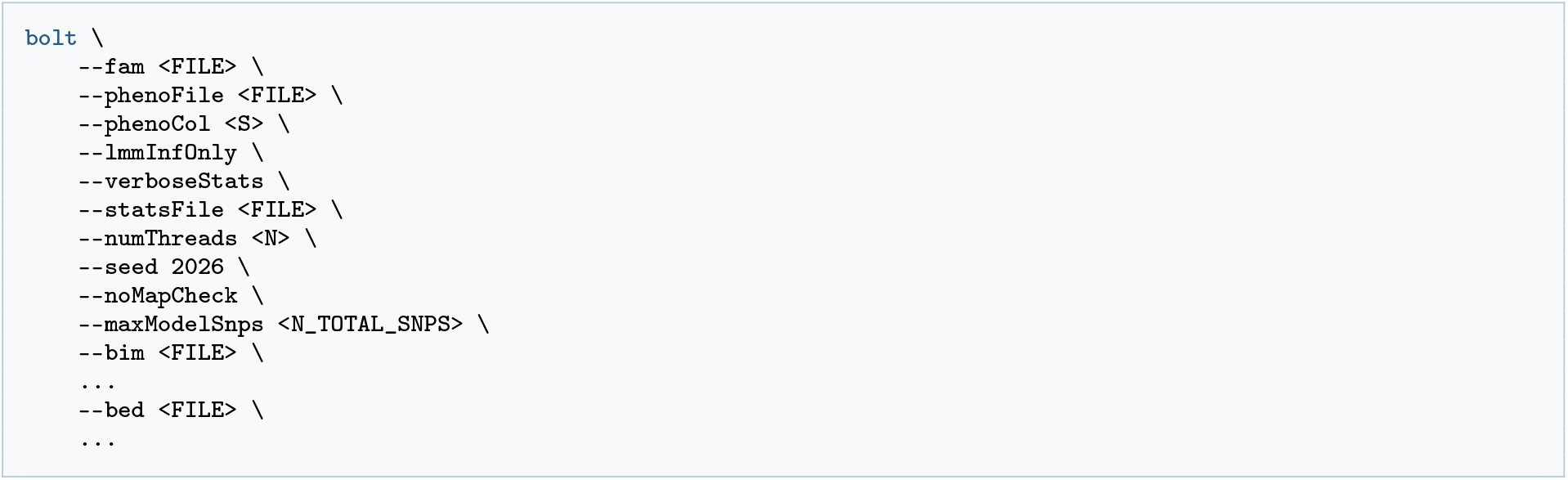

**Supplementary Table 13.** **BOLT-LMM** parameters.

| Application | Figure | --threads |
|---|---|---|
| BOLT-LMM | Fig. 4 | 64 |

### Supplementary Note 5 | GPU Memory Consumption and Requirement

Mikado GPU does not introduce significant additional memory overhead compared to the CPU version. GPU memory consumption comprises three components: the GRG, the input and output vectors, and the working memory.

1. The GRG occupies approximately the same space in GPU memory as it does on disk, plus the calculation buffer at each node;
2. The total size of the input and output vectors depends on the block width *k* and the data type;
3. The working memory can be classified into setup memory (for CUDA graphs, aliased value buffers, etc.) and application-specific memory.

The working memory remains well below the GRG size for the applications we evaluated. In Table 14, we provide the calculated sizes of (1) and (2), along with the measured maximum GPU memory usage across three different experiments.

**Supplementary Table 14.** GPU memory consumption.

| Application | Dataset | GRG Size | I/O Size | Total GRG + I/O | Measured Max. |
|---|---|---|---|---|---|
| GWAS | Sim 500k (Chr. 1) | 4.3 GB | 0.2 GB | 4.5 GB | 5.2 GB |
| PCA | 1000 Genomes | 11.9 GB | 0.5 GB | 12.4 GB | 13.3 GB |
| BOLT-LMM | 1000 Genomes Filtered | 6.3 GB | 0.8 GB | 7.1 GB | 9.0GB |

### Supplementary Note 6 | Platform Configuration

In Table 15, we list the hardware configurations of the machines used in this work.

**Supplementary Table 15.**
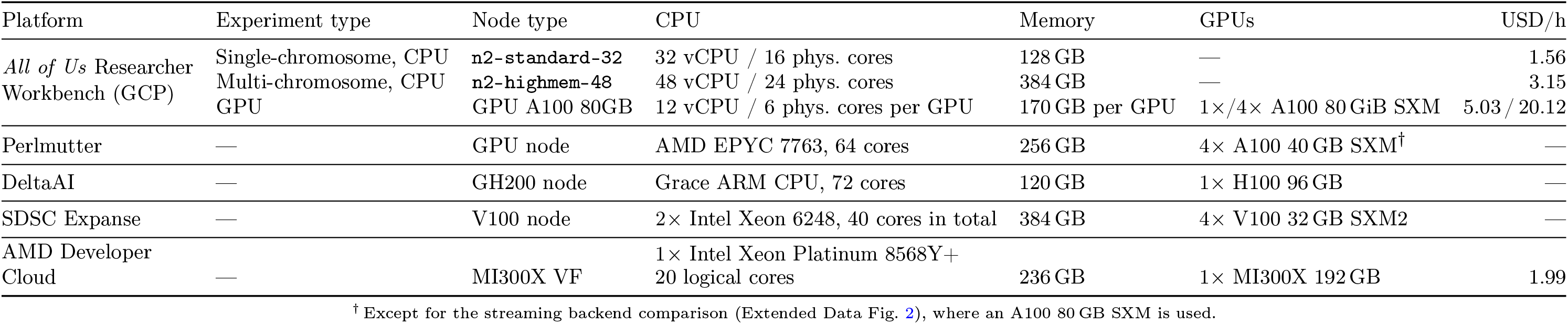
Platform configurations used in the experiments. V100 GPUs are connected by second-generation NVLink. A100 GPUs are connected by third-generation NVLink. GH200 nodes feature NVLINK-C2C. Cloud prices are based on rates at the time of writing.

## Footnotes

1 Mikado refers to the pick-up-sticks game, in which a stick may be lifted only when no other stick rests on it.

2 The factors are computed from unrounded timings and rounded for display.

3 https://github.com/CornellHPC/grapp/tree/c49cc2e37c0561cbd336b248b4091bb96782f897

4 Details on GRG construction can be found at https://grgl.readthedocs.io/en/stable/construct.html.

